# Heterochromatin-enriched assemblies reveal the sequence and organization of the *Drosophila melanogaster* Y chromosome

**DOI:** 10.1101/363101

**Authors:** Ching-Ho Chang, Amanda M. Larracuente

**Keywords:** *Drosophila melanogaster* genome, Y chromosome, long read assembly, gene duplications, gene conversion, *crystal-Stellate*

## Abstract

Heterochromatic regions of the genome are repeat-rich and gene poor, and are therefore underrepresented in even in the best genome assemblies. One of the most difficult regions of the genome to assemble are sex-limited chromosomes. The *Drosophila melanogaster* Y chromosome is entirely heterochromatic, yet has wide-ranging effects on male fertility, fitness, and genome-wide gene expression. The genetic basis of this phenotypic variation is difficult to study, in part because we do not know the detailed organization of the Y chromosome. To study Y chromosome organization in *D. melanogaster*, we develop an assembly strategy involving the *in silico* enrichment of heterochromatic long single-molecule reads and use these reads to create targeted *de novo* assemblies of heterochromatic sequences. We assigned contigs to the Y chromosome using Illumina reads to identify male-specific sequences. Our pipeline extends the *D. melanogaster* reference genome by 11.9-Mb, closes 43.8% of the gaps, and improves overall contiguity. The addition of 10.6 MB of Y-linked sequence permitted us to study the organization of repeats and genes along the Y chromosome. We detected a high rate of duplication to the pericentric regions of the Y chromosome from other regions in the genome. Most of these duplicated genes exist in multiple copies. We detail the evolutionary history of one sex-linked gene family—*crystal-Stellate*. While the Y chromosome does not undergo crossing over, we observed high gene conversion rates within and between members of the *crystal-Stellate* gene family, *Su(Ste)*, and *PCKR*, compared to genome-wide estimates. Our results suggest that gene conversion and gene duplication play an important role in the evolution of Y-linked genes.

## BACKGROUND

Heterochromatic regions of the genome are dense in repetitive elements and rarely undergo recombination *via* crossing over (Charlesworth *et al*. 1986). While heterochromatin is generally gene-poor, this compartment of the genome harbors functional elements (Gatti and Pimpinelli 1992) that impact a diverse array of processes, including nuclear organization (Csink and Henikoff 1996), chromosome pairing and segregation (Dernburg *et al*. 1996; McKee *et al*. 2000; Rosic *et al*. 2014), and speciation (*e.g*. Bayes and Malik 2009; Ferree and Barbash 2009; Cattani and Presgraves 2012). In many cases, the functionally relevant sequences are unknown, in part because it is difficult to sequence and assemble repeat-rich heterochromatic sequences. These sequences can be unstable in cloning vectors and/or toxic to *E. coli* cells (Carlson and Brutlag 1977; Lohe and Brutlag 1987b; Lohe and Brutlag 1987a) and thus may be underrepresented in clone-based sequencing libraries. Repetitive reads also present a challenge to most genome assemblers (Treangen and Salzberg 2011). As a result, many heterochromatic regions of the genome are missing from even the best genome assemblies (Hoskins *et al*. 2002; Carvalho *et al*. 2003). *Drosophila melanogaster* has arguably one of the most contiguous genome assemblies of any metazoan (Chakraborty *et al*. 2016). However, only ~143 Mb of the estimated ~180-Mb-genome is assembled into contigs (Hoskins *et al*. 2015). Heterochromatin makes up ~20% of the female and ~30% of the male *D. melanogaster* genome (the entire 40-Mb Y chromosome is heterochromatic; (Hoskins *et al*. 2002). The latest iteration of the reference genome assembly used BAC-based methods to extend into pericentromeric and telomeric regions, and increased the representation of the Y chromosome over 10-fold—the most recent genome assembly (version 6, R6 hereafter) includes ~27 Mb of heterochromatin, ~4 Mb of which is from the Y chromosome (Hoskins *et al*. 2015).

The Drosophila Y chromosome has been particularly recalcitrant to assembly (Hoskins *et al*. 2015). In addition to problems with cloning and assembly, we expect Y-linked sequences to have 50% and 25% of the autosomal coverage in male and mixed-sex sequencing libraries, respectively. Approximately 80% of the *D. melanogaster* Y chromosome is likely tandem repeats (Bonaccorsi and Lohe 1991). There are only ~20 known Y-linked genes (Carvalho *et al*. 2015), at least six of which are essential for male fertility (Kennison 1981). Despite being gene-poor, Y chromosomes can harbor functional variation. For example, structural variation on the Y chromosome in mammals affects male fertility (Reijo *et al*. 1995; Vogt *et al*. 1996; Sun *et al*. 2000; Repping *et al*. 2003). Similarly, Y-linked genetic variation in *D. melanogaster* has significant effects on male fertility (Chippindale and Rice 2001), including heat-induced male sterility (Rohmer *et al*. 2004). Y-linked variation in Drosophila also affects global gene expression (Lemos *et al*. 2008) and chromatin states across the genome (Lemos *et al*. 2010; Brown and Bachtrog 2014; Brown and Bachtrog 2017). It is unlikely that this functional variation maps to the few known Y-linked genes because there is very little nucleotide variation in coding regions (Zurovcova and Eanes 1999; Larracuente and Clark 2013). Instead, the Y chromosome may act as a sink for chromatin factors. Variation in the amount of Y-linked heterochromatin may influence the distribution of chromatin modifiers elsewhere in the genome (Dimitri and Pisano 1989; Henikoff 1996; Francisco and Lemos 2014; Brown and Bachtrog 2017). Without knowing the structure and composition of Y chromosomes, it is difficult to study this phenomenon in detail. Targeted attempts to sequence and assemble the Y chromosome have only had limited success (Hoskins *et al*. 2002; Abad *et al*. 2004; Mendez-Lago *et al*. 2009; Mendez-Lago *et al*. 2011; Hoskins *et al*. 2015; Mahajan *et al*. 2018). Single-molecule long read sequencing approaches (Branton *et al*. 2008; Eid *et al*. 2009) are improving our ability to assemble repetitive regions of complex genomes (Huddleston *et al*. 2014; Chaisson *et al*. 2015; Chang and Larracuente 2017; Jain *et al*. 2017; Khost *et al*. 2017) However, so far these approaches have only resolved relatively small segments of the Drosophila Y chromosome (Carvalho *et al*. 2015; Krsticevic *et al*. 2015).

Here we develop an approach using single-molecule long read sequencing from Pacific Biosciences (PacBio; Kim *et al*. 2014) to create heterochromatin-enriched genome assemblies. We use this approach to build a new assembly of the *D. melanogaster* genome that fixes gaps in euchromatin, adds a substantial amount of heterochromatin, and improves the overall contiguity of the genome assembly. Most of the additional sequence in our assembly is Y-linked, allowing us study Y chromosome composition in fine detail. We describe the landscape of transposable elements, the high rate of Y-linked gene duplication, and patterns of gene conversion among members of Y-linked multi-copy gene families.

## METHODS

### Heterochromatin sensitive assembly

We used BLASR (v5.1; Chaisson and Tesler 2012) to map PacBio reads (from Kim *et al*. 2014) to the release 6 (R6) *D. melanogaster* genome. Both the PacBio sequence reads and the reference genome are from the Iso1 strain. To curate a set of heterochromatin-enriched reads, we extracted any reads that: *1)* map outside of the major chromosome arms (i.e. 2L, 2R, 3L, 3R, 4, X) and mitochondria; or *2)* are unmapped. We took an iterative approach to genome assembly, generating two versions of both the heterochromatin and the whole genome assemblies, and then reconciling differences between them using quickmerge (Chakraborty *et al*. 2016). For the heterochromatin, we generated *de novo* assemblies with the heterochromatin-enriched reads using Canu v 1.3 (r7740 72c709ef9603fd91273eded19078a51b8e991929; Koren *et al*. 2017; repeat sensitive settings) and Falcon (v0.5; Chin *et al*. 2016; see Supplementary methods; Table S1). To improve the assembly of the major chromosome arms, we generated *de novo* assemblies with all PacBio reads using Falcon and Canu (Supplementary methods). We used quickmerge to combine our *de novo* heterochromatin-enriched assemblies with our all-read *de novo* assemblies sequentially, and then with two reference assemblies (R6; Hoskins *et al*. 2015) and a *de novo* PacBio assembly from (Chakraborty *et al*. 2016; Table S1). The detailed Falcon and Canu parameters for each *de novo* assembly and outline of the assembly and reconciliation process are in the supplementary methods. We also manually inspected each assembly, paying particular attention to Y-linked genes, where gaps in the assembly can occur because of low read coverage. We extracted raw or corrected reads from 7 Y-linked regions with read coverage < 10 and reassembled these manually in Geneious v 8.1.6 (Kearse *et al*. 2012). Before attempting to merge any assemblies, we checked that the gene order on all major chromosome arms agreed with R6 and examined the completeness of genes in pericentromeric regions, telomeres, and the Y chromosome. In our final reconciled assembly, we manually adjusted any errors in the *Rsp, Sdic*, and *Mst77Y* regions based on their organization in previous studies (Krsticevic *et al*. 2015; Clifton *et al*. 2017; Khost *et al*. 2017). We removed redundant contigs using MUMMER implemented in Masurca (v3.2.2; Zimin *et al*. 2017) and polished the resulting assembly using Quiver (SMRT Analysis v2.3.0; Chin *et al*. 2013). To correct any errors in regions with low PacBio coverage, we ran Pilon v1.22 (Walker *et al*. 2014) 10 times with both raw Illumina reads and synthetic reads (Table S2; with parameters “--mindepth 3 --minmq 10 --fix bases”). We created two and five scaffolds for the third and Y chromosomes respectively, based on known gene structure. We used MUMMER v3.23 (Kurtz *et al*. 2004) to map our new assembly to the R6 assembly using “nucmer --mum -l 10000 -D 40”, and only reported the one-to-one alignments using “delta-filter -1”. We remapped PacBio reads to this assembly using minimap v2.5-r607 (Li 2016) with parameters “-t 24 - ax map-pb”. We called coverage of uniquely mapped reads using samtools (v1.3 -Q 10; Li *et al*. 2009). To report on the sequence added in our assembly, we define heterochromatic regions based on the coordinates in Hoskins et al. (2015) and assume all added sequence beyond these coordinates on major chromosome arms, assigned to the Y chromosome, or on unassigned contigs is enriched in heterochromatin.

### Identifying Y-linked contigs

We used Illumina reads from male and female genomic libraries (Table S2) to identify Y-linked contigs. We mapped the male and female reads separately using BWA (v0.7.15; Li and Durbin 2010) with default settings and estimated the coverage of uniquely mapped reads per site with samtools (v1.3; -Q 10). We designated contigs with a median female-to-male read ratio of 0 as Y-linked (excluding sites with 1 or fewer Q>10 reads). To validate the sensitivity and specificity of our methods, we examined our X, Y, and autosome assignments for all 10-kb regions with a known location (only for regions with more than 1-kb of mappable sites).

### Gene and repeat annotation

We transferred r6.17 Flybase annotations from the R6 assembly to our final assembly using pBlat (v0.35, https://github.com/icebert/pblat-cluster; Kent 2002) and CrossMap (v0.2.5; Zhao *et al*. 2014). We then used HISAT2 (2.0.5; Kim *et al*. 2015) to map the male RNAseq reads (Table S2) to the genome based on known splice sites from the new annotation file. We used Stringtie (1.3.3b; Pertea *et al*. 2015) with these mapped reads and the guided annotation file from CrossMap to improve annotations and estimate expression levels. For unknown genes, we searched for homology using NCBI-BLAST against known *D. melanogaster* transcripts sequences (r6.17). To verify misassemblies and duplications, we designed primers to amplify segments of putatively Y-linked contigs/scaffolds with PCR in males and virgin females (Table S3). We also extracted and reverse transcribed RNA from 3-5 days old testes with TRIzol (ThermoFisher) and M-MLV reverse transcriptase (ThermoFisher), and examine splice sites using RT-PCR (Table S3).

To annotate repetitive DNA, we used RepeatMasker 4.06 (Smit *et al*. 2013) with Repbase 20150807 and parameters “-species drosophila -s”. We modified scripts from (Bailly-Bechet *et al*. 2014) to summarize TEs and other repetitive sequences. We searched for satellites using TRF (v4.09; Benson 1999) with parameters “2 7 7 80 10 100 2000 -ngs -h”.

### Sequence alignments and recombination analyses

We used BLAST v2.2.31+ (Altschul *et al*. 1990) and custom scripts to extract the transcript sequences from the genome. We aligned and manually inspected transcripts using Geneious v8.1.6 (Kearse *et al*. 2012). We constructed phylogenetic trees for regions conserved between members of the *cry-Stellate* family with MrBayes using the autosomal parent gene *Ssl* as an outgroup. (GTR+gamma HKY85 model; mcmc ngen=1,100,000 nchains=4 temp=0.2 samplefreq=200; seed=20,649). The consensus tree was generated with sumt burnin=500 with > 50% posterior probability. We used the APE phylogenetics package in R (Paradis *et al*. 2004) to plot the tree. We used compute 0.8.4 (Thornton 2003) to calculate Rmin and estimate population recombination rates based on linkage disequilibrium (Hudson 1987). In addition, we estimated gene conversion rates based on gene similarity (Supplementary methods; Ohta 1984; Rozen *et al*. 2003; Backstrom *et al*. 2005).

### Data availability

The genome assembly and annotations will be publicly available at Dryad. Supplemental materials (Figures S1-S5, Tables S1-S10, and File S1) are available at Figshare. The authors affirm that all data necessary for confirming the conclusions of the article are present within the article, figures, and tables.

## RESULTS

### Closing gaps in the Release 6 assembly

Major blocks of heterochromatin including the Y chromosome are missing from the latest version of the *D. melanogaster* genome (Hoskins *et al*. 2015). We built a new assembly of the *D. melanogaster* genome that closes gaps in the euchromatin and adds to the assembly in heterochromatin, most notably the Y chromosome. Even with long single molecule reads, unequal read coverage across heterochromatic regions may cause assembly problems (Carvalho *et al*. 2016). Because assemblers typically use the top ~30X longest reads for genome assembly, sex-linked regions may be under sampled. For example, some Y-linked regions are extremely underrepresented (e.g. there are no reads from the 3rd exon in *Ppr-Y* and only 9 reads come from the 2nd and 3rd exons of *kl-3)*. To reduce this potential bias, we assembled the heterochromatin and euchromatin separately and then combine these assemblies with each other and with published versions of the *D. melanogaster* genome (Fig. 1). We first isolate a set of heterochromatin-enriched reads by mapping all Pacbio reads to the R6 reference and discarding reads mapping uniquely to the euchromatic genome (Fig 1A). Using this approach, we extracted ~1.58 Gb of sequence across 204,065 reads (12% of total reads) for reassembly. With this small subset of reads, we are able to optimize parameters for repeat assembly, partially remedy assembly errors, and increase assembly contiguity. For Canu, we experimented with assembly conditions by varying bogart parameters (see Supplementary methods). For Falcon, we experimented with the minimal overlap length in the string graph. For both methods, we identified parameter combinations that maximized assembly N50, total assembly length, and longest contig length; and without detectable misassemblies in Y-linked coding regions. We note that while assembly length and contiguity are often used to assess assembly quality, the most contiguous assemblies are not always correct (Khost *et al*. 2017). We therefore reconciled the assembled contigs from the two “best” versions of our heterochromatin-enriched and whole genome assemblies sequentially, and finally with the R6 assembly and another PacBio reference assembly (Fig 1B-C; Chakraborty *et al*. 2016). We manually adjusted misassembled contigs and polished the final assembly for use in downstream analyses (Fig 1D). Our final reconciled genome has 200 contigs and is 155.6 Mb in total—this is a great improvement in assembly contiguity over R6 (143 Mb in 2,442 contigs; Table 1). The improvement is in both euchromatic and heterochromatic regions (Figs S2 and S3).

**Figure 1.**
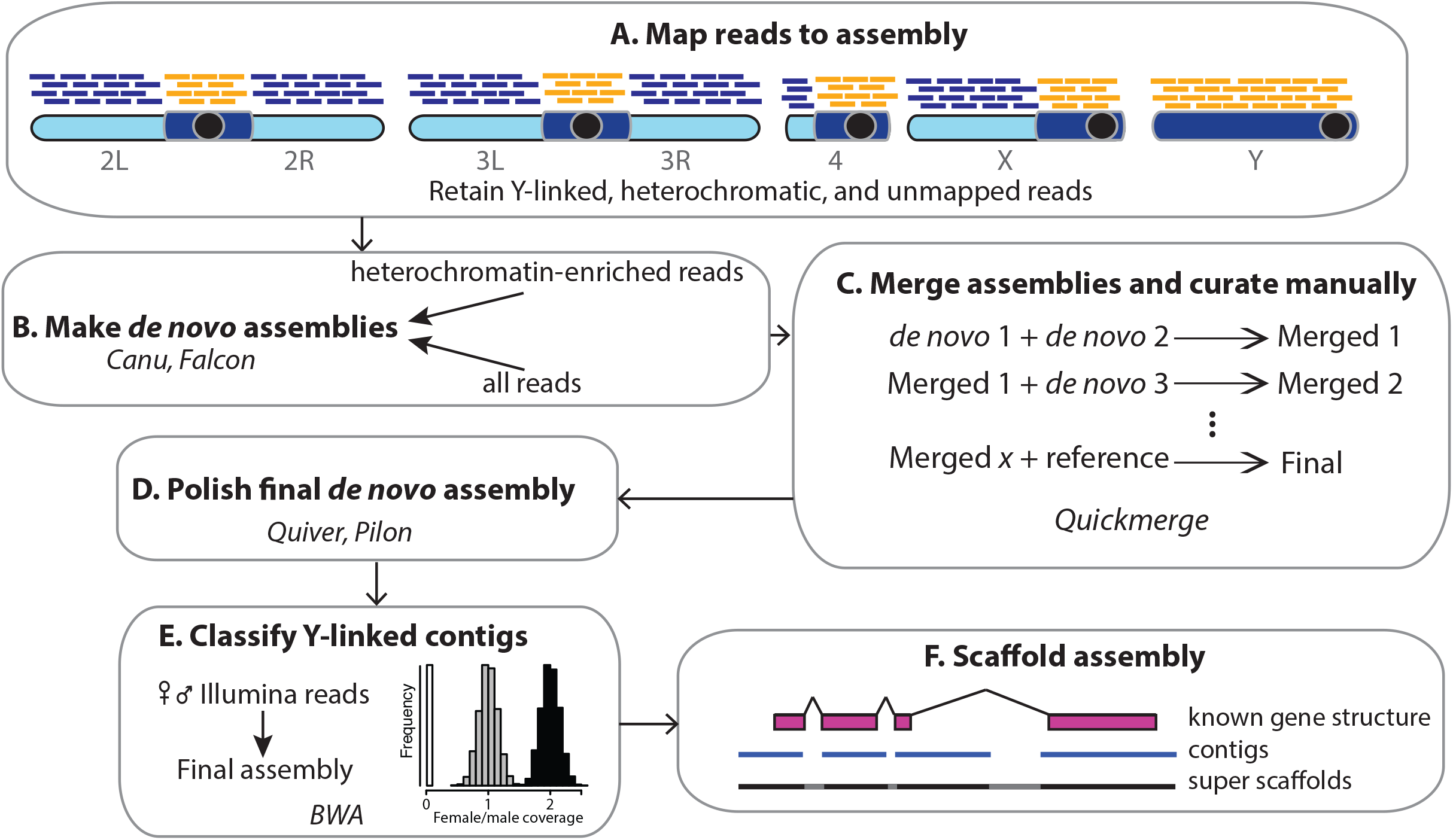
Overview of the heterochromatin-enriched assembly approach. **A)** We obtain a set of heterochromatin-enriched PacBio reads by mapping reads to the R6 assembly and retaining reads that map to known pericentric heterochromatin, Y chromosome contigs, or are unmapped (orange lines). **B)** We generate separate *de novo* PacBio assemblies for all reads (orange+blue lines) and for heterochromatin-enriched reads (orange lines) with Canu and Falcon. **C)** We merge assemblies sequentially using quickmerge to create the final assembly (Table S1). All assemblies were manually inspected and adjusted (see Methods). **D)** We polished the final *de novo* assembly with one round of quiver (using raw PacBio reads) and 10 iterations of Pilon (using male Illumina reads). **E)** We assign contigs in the final assembly to the X, Y, or autosomes using relative mapping of female-to-male Illumina reads (see Methods). **F)** Finally, we join contigs into super scaffolds using exon orientation information from known gene structures.

**Table 1.** Heterochromatin-enriched *D. melanogaster* assembly statistics

| Assemblies |  | Summaries |  |
| --- | --- | --- | --- |
| Whole Genome | # contigs | Total size | Contig N50 |
| GCF_000001215.4 (R6) | 2442 | 143,726,002 | 21,485,538 |
| This study | 200 | 155,584,520 | 21,691,270 |
| Y chromosome |  |  |  |
| GCF_000001215.4 (R6) | 261 | 3,977,036 | 81,922 |
| This study | 80 | 14,578,684 | 416,887 |

Our new assembly fills all three unassembled gaps in the gene-rich regions of the R6 major chromosome arms (one each on 2R, 3L and 4; Fig S2 and Table S5). Chromosome 4 had a predicted 17-kb gap in R6. In agreement with this predicted gap size, our new assembly inserts 17.996 kb in this gap with (AAATTAT)_n_ repeats and other AT-rich sequences. The gap on chromosome 2R was unsized; our assembly fills this gap with 4,664 bp consisting of 123-bp complex repeats. Interestingly, an annotated non-coding gene, CR44666, is located near the 2R gap in R6 and consists entirely of this 123-bp unit. In agreement with the predicted gap size of ~7kb on 3L, our new assembly inserts 6,157 bp containing one of four tandem copies of the 3S18/BEL transposons. Our assembly therefore places all euchromatic regions of the major chromosome arms on single contigs.

We also made a marked improvement to heterochromatic regions (as defined by Hoskins *et al*. 2015). In total, we filled 25 of 57 gaps in the R6 major chromosome scaffolds (Table S5). Of these gaps, 14 were located in transposon dense regions; four were associated with complex repeats (two with *Responder*, one with *1.688* family repeats and one with a newly-identified 123-bp unit), three were associated with 7-bp tandem repeats, and one is associated with rDNA repeats. One is a 17-kb deletion and the other two gaps involve complex rearrangements between R6 and our assembly that may represent scaffolding errors in R6. Our new assembly has ~38.6 Mb of heterochromatin-enriched DNA across 193 contigs, whereas the R6 assembly has ~26.7 Mb of heterochromatin-enriched DNA in 2,432 contigs. Approximately 89% of the additional heterochromatic sequence in this assembly is from the Y chromosome (see below). We assigned some contigs based on their repeat content, *e.g*. a 180-kb contig from chromosome *2* (Contig 142). This contig terminates in (AATAACATAG)_n_ and (AAGAG)_n_ repeats mapping to cytological bands *h37* and *h38* (Garavis *et al*. 2015). Contig 142 extended an existing unmapped R6 scaffold (Unmapped_Scaffold_8_D1580_D1567), which contains a gene (*klhl10)* that maps to chromosome 2 (http://flybase.org/reports/FBgn0040038).

### Identifying Y-linked contigs

The estimated size of the Y chromosome is 40 Mb, however only ~4 Mb is assembled and assigned to the Y chromosome in R6 (Hoskins *et al*. 2015). Our assembly pipeline based on PacBio reads circumvents the cloning steps associated with BAC-based sequencing, and results in a better representation of heterochromatin, including the Y chromosome. We developed an approach to identify and assign Y-linked contigs based on detecting male-specific sites using Illumina reads (Fig 1E). To validate our method to assign Y-linkage, we used contigs with a known location in R6 as benchmarks. Previous studies in mosquitos and *D. melanogaster* identified Y-linked contigs using the chromosome quotient (CQ): the female-to-male ratio of the number of alignments to a reference sequence (Hall *et al*. 2013). In *D. melanogaster*, this method has 76.3% sensitivity and 98.2% specificity (Hall *et al*. 2013). Our approach instead considers the number of male-specific regions (where the median per-site female-to-male ratio = 0) and is a better indicator of Y-linkage than CQ—among 14,116 10-kb regions in our assembly with known chromosomal location based on previous data (R6 assembly), we appropriately assigned 99.0% of Y-linked regions (714/721 regions; Fig S4). Only 1.5% of all regions that we assigned to the Y chromosome are not Y-linked in the R6 assembly (11/725 regions; Fig S4). Therefore, our method has both a higher sensitivity and specificity than previous methods. For the 11, 10-kb regions that may be false positives in our method, 9 are from a centromeric scaffold (3Cen_31_D1643_D1653_D1791), and 2 are from the 2^nd^ chromosome telomeres. These regions may be misassigned in the R6 assembly because the centromeric scaffold has a Y-specific repeat, AAAT, (Wei *et al*. 2018) and telomeric transposons are found on all chromosomes and may vary within strains. We used our method to assign 14.6 Mb to the Y chromosome across 106 contigs (N50 = 415 kb; Table 1). Because ~80% of the 40-Mb Y chromosome consists of tandem repeats (Lohe *et al*. 1993), this is likely near the maximum amount of Y-linked sequence we can expect to identify with current sequencing technology.

### Improving known Y-linked gene annotations

The gene order and orientation of Y-linked genes in our assembly is consistent with previous mapping data (Fig. 2; Carvalho *et al*. 2000; Carvalho *et al*. 2001; Vibranovski *et al*. 2008) using Y chromosome deletions, except for *Pp1-Y1*. Unfortunately, we cannot distinguish whether this difference is due to a misassembly or strain variation. We found splice site errors in two previous Y-linked gene models: *kl*-5 has three additional introns (in the 1^st^, 5^th^ and 17^th^ exons of the R6 annotation; Table S6), and *CCY* has one additional intron (in the 6^th^ exon of the R6 annotation; Table S6). We also found partial duplications of exons in *kl-3, ORY, PPr-Y*, and *WDY* (Table S7). Each of these duplications, except *ORY*, exists on unannotated regions of the R6 assembly. In the R6 assembly, *CCY* and *kl-3* contain misassembled sequences in 6th and 3rd exons coding region, respectively. We therefore corrected the misassemblies in the R6 Y-linked coding regions based on our assembly and PCR validation (Table S3). We used gene annotation data to scaffold the assembly (Fig 1F).

**Figure 2.**
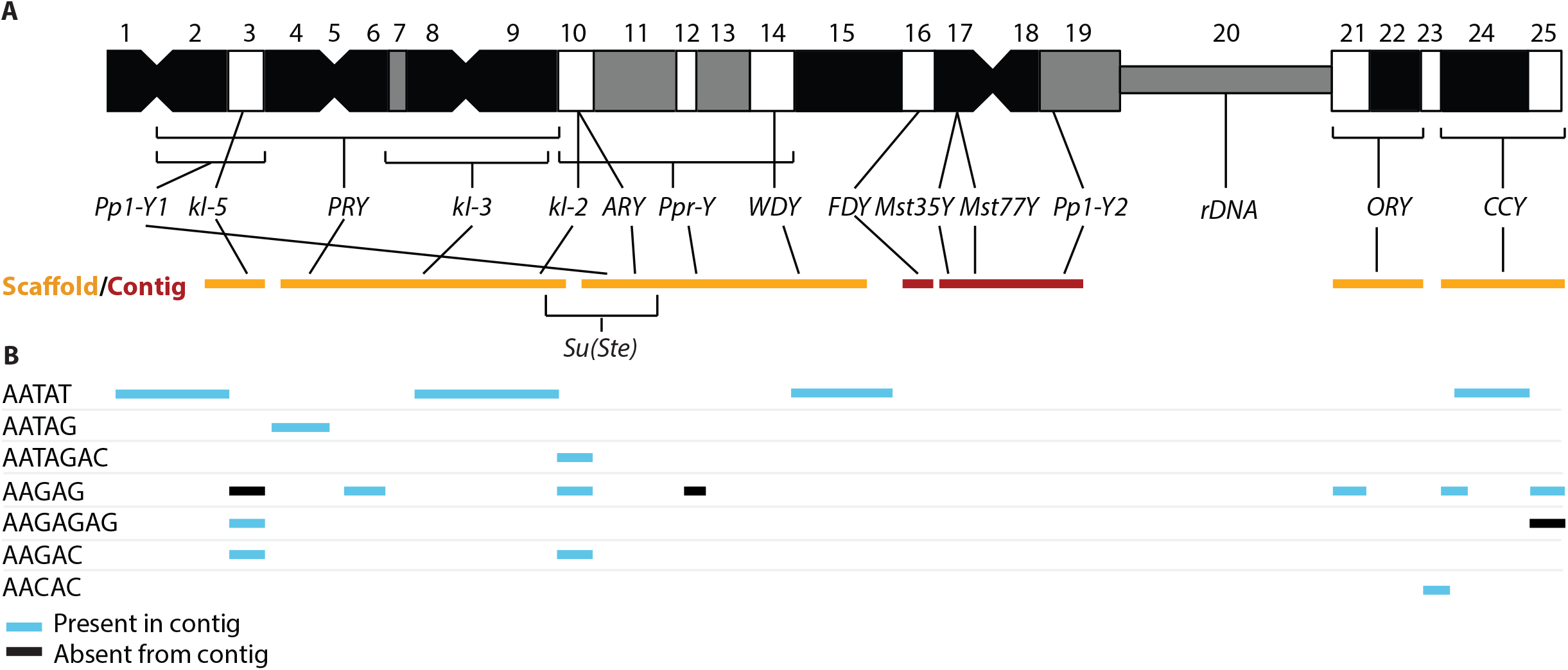
Schematic of Y chromosome organization. **A)** The Y chromosome is organized into 25 cytological bands. The position of the Y-linked genes is shown based on deletion mapping (Carvalho *et al*. 2000; Carvalho *et al*. 2001; Vibranovski *et al*. 2008). The major scaffolds (orange bars) and contigs (dark red bars) that span each Y-linked gene, from left to right are: Y_scaffold6, Y_scaffold7, Y_scaffold4, Y_Contig10, Y_Contig2, Y_scaffold5, and Y_scaffold3. Note that scaffolds may still contain gaps. **B)**’ The approximate cytological location of large blocks of simple tandem repeats (Bonaccorsi and Lohe 1991) agrees with the organization of our scaffolds and contigs: blue bars indicate that a block of satellite appears in that contig/scaffold, and black bars indicate that a block of repeats is missing from that contig/scaffold. Note that missing repeats may fall entirely in the gaps in our scaffolds, and potential cross-hybridization between AAGAG and AAGAGAG might explain the three discrepancies between our assembly and the cytological map.

### Y-linked gene duplications

We identified 13 independent duplications to the Y chromosome from other chromosomes, seven of which we identify as Y-linked for the first time. 11 of these duplications exist in multiple copies on the Y chromosome (Table 2). We also identified a new Y-linked gene, *CG41561* located on an unmapped contig (211000022280328) in the R6 assembly (Mahajan and Bachtrog 2017). Among the 13 duplications, we found that the Y-linked copies of *Hsp83, Mst77F (Mst77Y)*, and *vig2 (FDY)* are still expressed in testes (FPKM > 5 in at least one dataset; Table S8); however, *Hsp83* contains a premature stop codon and a TE insertion. Therefore, outside of *Mst77Y* and *FDY*, we do not have evidence for their function (Krsticevic *et al*. 2010; Krsticevic *et al*. 2015). Interestingly, these duplications seem to be clustered on the Y chromosome: six of duplications are on Y_scaffold4 and five of the duplications are on Y_Contig2 (Table 2). Y_scaffold4 and Y_Contig2 are from the cytological divisions *h10-15* and *h17-18*, respectively (Fig. 2). Additionally, *FDY* (Y_Contig10) maps to *h15-h20* (Krsticevic *et al*. 2015). Therefore, 12 of the duplications are located between *h10-h20* (11 of 25 Y-linked cytological bands), suggesting that the pericentromere of the Y chromosome (defined here as *h10-h20*) is enriched for duplicated genes in *D. melanogaster* (Fisher’s exact test; *P* = 0.005).

**Table 2.** Translocations to the Y chromosome from the autosomes and X chromosome

| Parent genes | Parent | Y copy # | Location of duplication on Y | source | name | Citation |
| --- | --- | --- | --- | --- | --- | --- |
| *Gs1l* | 2L | 2 | Y_scaffold4 | DNA | NA | Tobler et al. 2017 |
| *smt3* | 2L | 5 | Y_scaffold4, Y_Contig140,<br>Y_Contig23 | RNA | NA | NA |
| *ProtA* | 2L | 9 | Y_Contig2, Y_Contig6, Y_Contig104 | DNA | *Mst35Y* | Méndez-Lago et al. 2011 |
| *Hsp83* | 3L | 6 | Y_scaffold4 | RNA | NA | NA |
| *velo* | 3L | 70 | Y_Contig2, Y_Contig6, Y_Contig104 | Unknown | NA | NA |
| *Pka-R1, CG3618, Mst77F* | 3L | 15,17,18 | Y_Contig2 | DNA | *Mst77Y* | Krsticevic et al. 2010 |
| *Dbp80* | 3L | 1 | Y_scaffold6 | DNA | NA | NA |
| *fru* | 3R | 6 | Y_scaffold4 | Unknown | NA | NA |
| *CG5886* | 3R | 2 | Y_scaffold4 | Unknown | NA | NA |
| *vig2, Mocs2, Clbn, Bili* | 3R | 1,1,7,1 | Y_Contig10 | DNA | *FDY* | Carvalho et al. 2015 |
| *Tctp* | 3R | 1 | Y_scaffold4 | Unknown | NA | NA |
| *CR43975* | 3R | 78 | Y_Contig2, Y_Contig4, Y_Contig6,<br>Y_Contig104, Y_Contig22 | DNA | NA | Tobler et al. 2017 |
| *CG12717, ade5* | X | 214,33 | Y_Contig2, Y_Contig6, Y_Contig104 | DNA | NA | Méndez-Lago et al. 2011 |
| *CG41561* | U* | 1 | Y_Contig74 | NA | NA | Mahajan and Bachtrog 2017 |

\* Unmapped contig 211000022280328

### Repeat content in Y-linked contigs

Cytological observations indicate that the Y chromosome is highly enriched for repetitive sequences (Lohe *et al*. 1993; Carmena and Gonzalez 1995; Pimpinelli *et al*. 1995), however there have not been attempts to document this at the sequence level. We used our assembly to identify repetitive elements across the Y chromosome. Consistent with previous studies, we find that the Y chromosome is enriched for rDNA and IGS repeats (Ritossa and Spiegelman 1965; Fig 3A and Table S9). The rDNA are located across 54 scaffolds/contigs, including 1 Y-linked scaffold, 12 Y-linked contigs, 2 X-linked contigs, and 39 unknown contigs (Table S9). We identified 56 copies of 18s rDNA, 238 copies of 28s rDNA, and 721 copies of IGS repeats on the Y chromosome. LTR and LINE transposons contribute 53% and 19% of the total sequence, respectively, in our Y-linked contigs (Fig 3A). We assume that most of the unassembled parts of the Y chromosome are simple tandem repeats (Lohe *et al*. 1993). Based on this assumption, we estimate that 65% of the 40-Mb Y chromosome is simple tandem repeats, and LTR and LINE elements comprise 18% and 7% of the total 40-Mb Y chromosome, respectively. Compared to the rest of the genome, the Y chromosome has a 1.4 - 1.8 fold enrichment of retrotransposons (10.2% of LTR and 5.0% of LINE for the rest of the genome), while DNA transposon content is similar among chromosomes (2.3% on Y and 2.2% for the rest of the genome, Fig. 3A). The Y chromosome is enriched for retrotransposons over DNA transposons even when compared to other heterochromatic genomic regions (Fig. S5).

**Figure 3.**
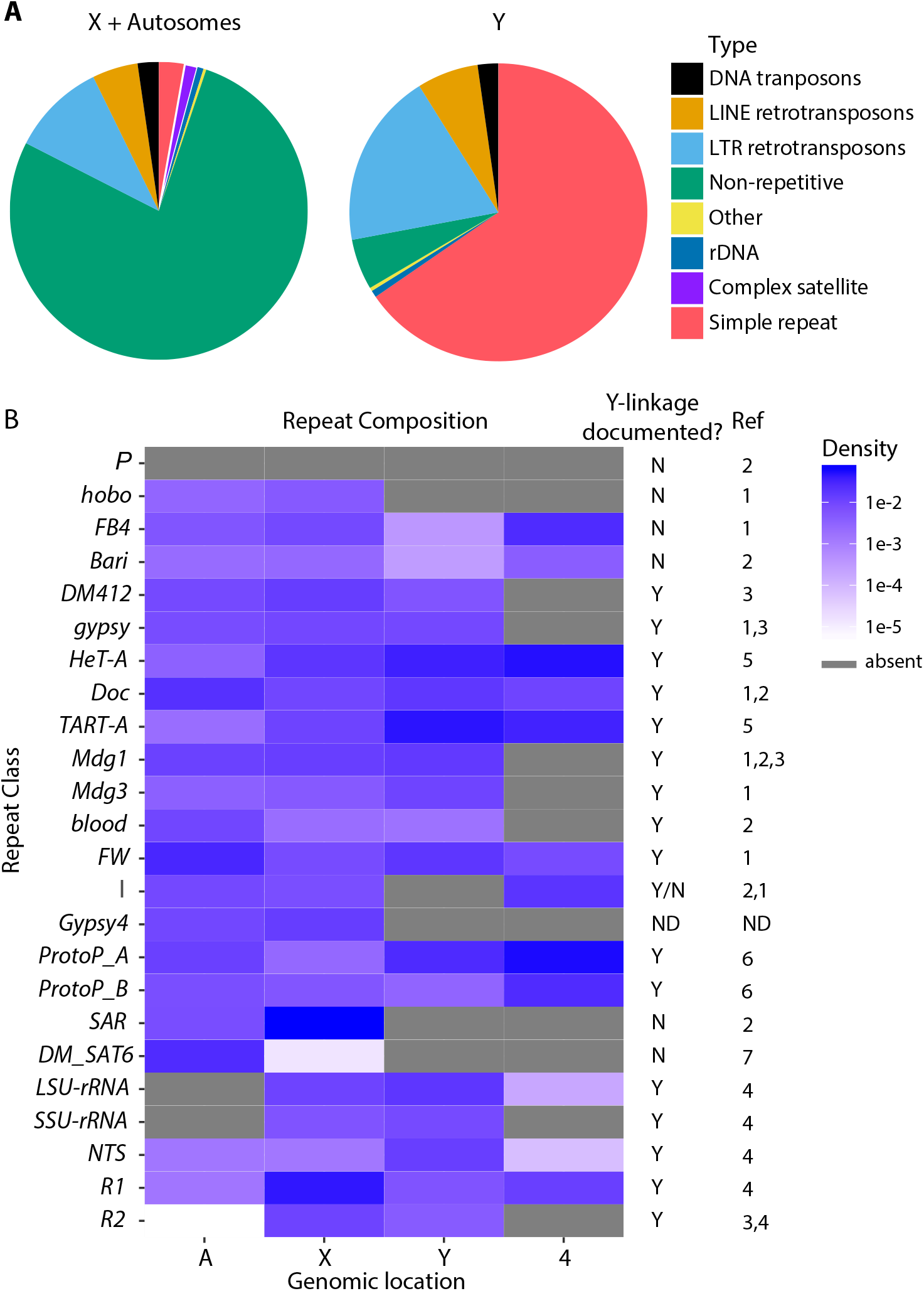
Repeat composition on the Y chromosome compared to the rest of the genome. **A)** The major repeat class composition on Y-linked contigs and all other contigs in our assembly (from the X and autosomes). **B)** A comparison of complex repeats and transposable elements between autosomes, X, Y, and 4th chromosomes. We indicate the presence/absence (Y/N, respectively) of repeat classes for which cytological and/or Southern hybridization data exists in the literature. I-elements have conflicting reports of Y-linkage in the literature. References: 1.Carmena and Gonzalez 1995; 2. Pimpinelli *et al*. 1995; 3.Junakovic *et al*. 1998; 4. Ritossa and Spiegelman 1965; 5. Agudo *et al*. 1999; 6. Balakireva *et al*. 1992; 7. Abad *et al*. 1992.

Previous studies predicted the repeat composition of the Y chromosome based on the presence/absence of *in situ* hybridization (ISH) signals on mitotic chromosomes. (Carmena and Gonzalez 1995; Pimpinelli *et al*. 1995). Our assemblies recapitulate these ISH results. For example: P, *hobo, FB4*, and *Bari-1* are nearly absent from the Y chromosome (< 3.5 kb of total sequence), while *Dm412, Gypsy, HetA, Doc, TART, Mdg1, Mdg3, blood*, and *FW* have at least 14 kb of sequence on the Y chromosome (Fig 3B and Table S9; Carmena and Gonzalez 1995; Pimpinelli *et al*. 1995; Junakovic *et al*. 1998; Agudo *et al*. 1999). Previous studies are conflicted about the presence/absence of I elements (Carmena and Gonzalez 1995; Pimpinelli *et al*. 1995), however we do not see evidence of Y linkage in our assembly. Other transposons also appear to be absent from the Y chromosome, *e.g. gypsy4* (Table S9; Fig 3B). Since *I* element-mediated dysgenesis only occurs in females (Bucheton *et al*. 1976), it is possible that this element is inactive in the male germline and therefore rarely has the opportunity to invade Y chromosomes. We suggest that the sex-specific activity of TEs may contribute to their genomic distribution.

Tandem repeats are also enriched on Y chromosomes (~65% on the Y chromosome compared to 2.8% on the other chromosomes; (Lohe *et al*. 1993). Approximately 5% (742,964 bp) of our Y-linked sequences are tandem repeats. We assume that this is a gross underestimate of tandem repeat abundance, but nevertheless helps lend insight into the repeat content and organization of the Y chromosome. Our assembly agrees with most previous cytological and molecular evidence of Y chromosome simple tandem repeat content (Fig. 2; Bonaccorsi and Lohe 1991). Among 32 known Y-linked simple repeats, 20 appear in our Y-linked contigs (Table S10; Bonaccorsi and Lohe 1991; Jagannathan *et al*. 2017; Wei *et al*. 2018). The repeats that we do not find may be sequence variants of abundant repeats (e.g. we detect AAAAC and AAAGAC but not AAAAAC or AAAAGAC), not perfectly in tandem, or part of a more complex repeat (e.g. AAGACAAGGAC is part of AAGACAAGGAAGACAAGGACAAGACAAGGAC; Table S10). Although we recover only ~60% of known Y-linked repeats (based on Illumina data, Wei *et al*. 2018; or ISH, Bonaccorsi and Lohe 1991; Jagannathan *et al*. 2017), our new assembly including genes and transposable elements provides the most detailed view of Y chromosome organization.

### Evolution of the *cry-Stellate* gene family

The multicopy *crystal-Stellate (cry-Ste)* gene family is thought of as a relic of intragenomic conflict between X and Y chromosomes (reviewed in Bozzetti *et al*. 1995; Hurst 1996; Malone *et al*. 2015). *Stellate* (Ste) is an X-linked multicopy gene family whose expression is controlled by the Y-linked *Suppressor of Stellate* (Su(Ste)) locus through an RNA interference mechanism (Nishida *et al*. 2007). If left unsuppressed, *Ste* expression leads to the accumulation of crystals in primary spermatocytes of the testes, resulting in male sterility (Bozzetti *et al*. 1995). This multicopy gene family has a complicated evolutionary history (Kogan *et al*. 2000). *Ste* and *Su(Ste)* are recent duplications of the autosomal gene *Su(Ste)-like (Ssl* or *CK2β*) with a testis-specific promoter from casein kinase subunit 2 (Kogan *et al*. 2000). Following the initial duplication of *Ssl* to the Y chromosome, members of this gene family expanded and duplicated to the X chromosome (Fig 4A). All sex-linked members of this gene family exist in multiple copies. The X-linked copies and Y-linked copies amplified independently, perhaps driven by sex chromosome conflict (Kogan *et al*. 2012). We used our assembly to study the evolution of this interesting gene family and patterns of gene conversion on the Y chromosome. We found 666 copies of genes in the *cry-Ste* family: 37 on the X chromosome, 627 on the Y chromosome, and 2 from an unknown region. We detect more Y-linked copies than were previously estimated (200-250 complete copies) using southern blotting (McKee and Satter 1996). We found a clade of 122 Y-linked genes that are from an ancestral duplication of *Ssl* and fall as an outgroup to *Ste* and *Su(Ste)* (Fig. 4B). These copies, originally identified in a Y-derived BAC, are designated as pseudo-CK2β repeats on the Y chromosome (*PCKRs)* and have the ancestral promoters (Danilevskaya *et al*. 1991; Usakin *et al*. 2005). However, there is very little expression among the 107 copies of *PCKR* (total FPKM <3 from FBtr0302352 and MSTRG.17120.1; Table S8). *Ste* copies appear in both the X heterochromatin and euchromatin (hereafter referred to as *hetSte* and *euSte*, respectively; (Livak 1984; Shevelyov 1992). In addition to the 13 previously-assembled copies of *euSte* (cytological divisions 12E1 to 12E2), we found an additional 20 copies of *Ste* located on two X-linked contigs (17 on Contig5 and 3 on X_9), corresponding to functional *hetSte* copies, and pseudogenized tandemly-repeated heterochromatic elements related to *Stellate* (SCLRs; Nurminsky *et al*. 1994; Tulin *et al*. 1997). The 3 SCLRs on the contig X_9 were present but not annotated in the R6 assembly and were located proximal to *hetSte*. We assembled 17 *hetSte* in a single 500-kb contig, where two *hetSte* loci (5 and 12) are separated by *BATUMI* and rDNA sequences. However, previously published data using restriction maps and southern blotting suggests that *hetSte* are organized into three loci (with ~14, 3, and 4 copies) separated by *BATUMI* and rDNA (Tulin *et al*. 1997). Our phylogenetic analysis reveals that SCLRs and *hetSte* are clustered, suggesting that *hetSte* and *euSte* amplified independently or experience concerted evolution (Fig. 4B).

**Figure 4.**
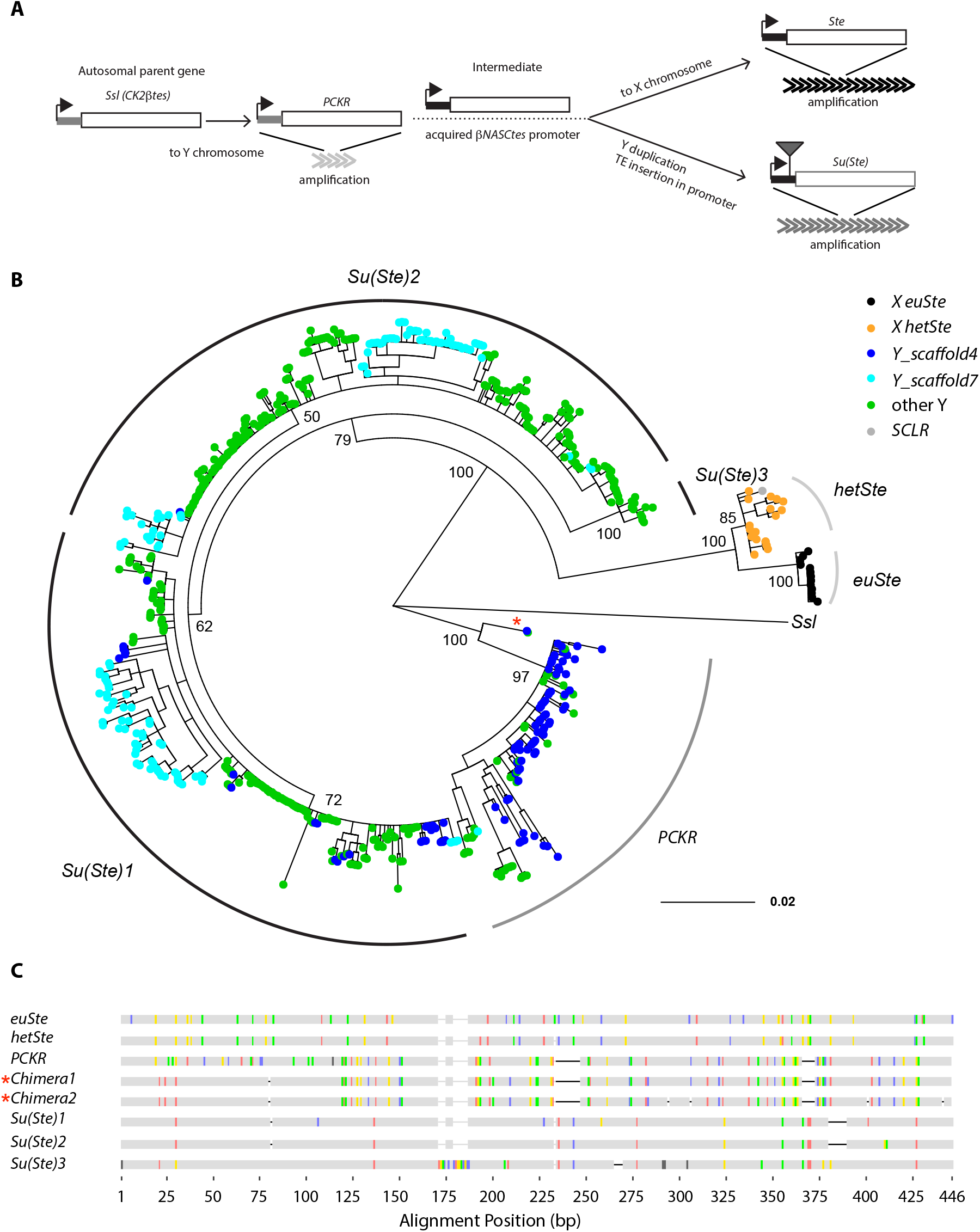
Evolution of the *Cry-Ste* family. **A)** The evolutionary history of *Cry-Ste* family in *D. melanogaster* (modified from Usakin *et al*. 2005); **B)** A Bayesian phylogenetic tree constructed with 606 full-length copies of genes in the *Cry-Ste* family including *Ssl* (parent gene) as the outgroup. Tip colors represent the location of genes in our assembly. Posterior node confidence is shown for a subset of the primary nodes separating repeat types. SCLR is a non-functional variant of *Ste*. **C)** The alignment of representative repeats for heterochromatic Ste (*hetSte)*, euchromatic Ste (*euSte*), *PCKR*, three main variants of *Su(Ste*), and two chimeric genes are shown (also indicated with red * in tree). Vertical colored lines indicate where base changes (red=A; yellow=G; green=T; blue=C; grey=missing) occur and dashes indicate indels.

The 627 *Su(Ste)* and *PCKR* copies are spread across 10 and 3 Y-linked contigs, respectively. These repeats primarily occur in tandem and are flanked by different transposon sequences, including *1360, Gypsy12*, and the telomere-associated transposons, *HeT-A, TART*, and *TAHRE*. Previous studies suggested that the acquisition of *1360* in *Su(Ste)* may have been an important step in *Su(Ste)* evolving a piRNA function to suppress *Ste* (Usakin *et al*. 2005). *HeT-A* colocalizes with Ste-like sequences in the BAC Dm665 (Danilevskaya *et al*. 1991). We found that the Ste-like sequences in Dm665 are PCKRs and are located proximal to *Su(Ste)*, between *WDY* and *Pp1-Y1*. Consistent with BAC data and our assembly, this region is also enriched for telomeric sequences (based on ISH, Traverse and Pardue 1989; Abad *et al*. 2004). Interestingly, we found 2 chimeric copies of *PCKR* and *Su(Ste)* (Fig. 4C), suggesting inter-genic gene conversion occurred between these genes. Previous studies hypothesized that gene conversion homogenizes *Su(Ste)* clusters, but these studies were only based on restriction maps or a few variants (Balakireva *et al*. 1992; McKee and Satter 1996). We investigated the rate of gene conversion on the Y chromosome using 107 copies of *PCKR* and 406 copies of *Su(Ste)* after removing fragments smaller than 280 bp. We detected evidence of recombination at both *PCKR* (per 857-bp locus: Rmin=2 and *ρ* = 2.67; *c_g_* = 2.9 x 10^-5^ events per site, per generation) and Su(Ste) (per 1203-bp locus: Rmin=1 and *ρ* = 4.04; *c_g_* = 8.2 x 10^-6^ events per site, per generation). Since there is no recombination *via* crossing over, we estimate the Y-linked gene conversion rate to be 0.8-5 x 10^-5^ events per site, per generation. We also used estimates of similarity among repeats within each gene family to estimate gene conversion rates (Supplementary methods; *c_g_*). Assuming a mutation rate of 2.8 x 10^-9^ per site per generation (Keightley *et al*. 2014) we estimate the rate of gene conversions per site per generation to be 2.1 x 10^-5^ and 1.5 x 10^-4^ for *PCKR* and *Ste*, respectively. These rates are ~10^3^ – 10^4^ times higher than gene conversion rates on the autosomes and X chromosome (Comeron *et al*. 2012; Miller *et al*. 2012; Miller *et al*. 2016), and surprisingly similar to the rate observed in mammalian Y and bird W chromosomes (Repping *et al*. 2003; Backstrom *et al*. 2005); both based on *c_g_*). Rmin and LD-based estimators may underestimate the true gene conversion rate because both recent amplification and selection could decrease variation among copies and cause us to miss recombination events. On the other hand, we likely overestimate the gene conversion rate based on similarity among copies for the same reasons. With both approaches, our data suggest high rates of intrachromosomal gene conversion on Y chromosomes. Recombination may also occur between the X and Y chromosomes—of the 116 variant sites in *Ste*, 62 of the same variants are found at the homologous positions in *PCKR* and/or *Su(Ste)*. It will be important to further explore rates of Y-linked gene conversion using multiple strains of *D. melanogaster*. Higher gene conversion rates in Y-linked multicopy gene families may be important for the evolution of Y-linked genes.

## Discussion

Heterochromatic sequences can contain important genetic elements (e.g. Gatti and Pimpinelli 1992) but tend to be underrepresented in genome assemblies. Single-molecule real time sequencing is making strides towards achieving complete assemblies of complex genomes (Huddleston *et al*. 2014; Chaisson *et al*. 2015), however densely-repetitive regions still present a significant assembly challenge that often requires manual curation (Krsticevic *et al*. 2015; Clifton *et al*. 2017; Khost *et al*. 2017). Uneven read coverage across the genome, and lower read coverage in heterochromatic regions, likely cause problems with genome assembly (Krsticevic *et al*. 2015; Chang and Larracuente 2017; Khost *et al*. 2017). Our assembly approach is based on the *in silico* enrichment of heterochromatic reads, followed by the targeted reassembly of heterochromatic regions, and finally, a reconciliation between whole genome and heterochromatin-enriched assemblies. This approach helped fill gaps, fix errors, and expand the *D. melanogaster* reference assembly by 11.9 Mb (8% more sequence than the latest release, R6).

Approximately 89% of the additional sequence in our assembly is from the Y chromosome, allowing us to get a detailed view of Y chromosome organization. Despite these improvements, we are still missing some Y-linked regions and some required manual correction. Assemblers filter reads when they appear chimeric or where pairs of reads disagree about overlaps. Canu and Falcon tend to disagree about the organization of some highly repetitive sequences (e.g. *Rsp*, Khost *et al*. 2017; *Sdic*, Clifton *et al*. 2017; and *Mst77Y*, Krsticevic *et al*. 2015). Our approach does not completely remedy this problem—we also identified errors in our preliminary assemblies that required manual correction. For these misassembled regions, Falcon and Canu arrive at different sequence configurations (e.g. we found 20 copies of *Mst77Y* in the Canu assembly and 14 copies in the Falcon assembly). To resolve these differences, we leveraged evidence from ISH studies and known gene structures to identify and reconcile differences between the assemblies. Our results suggest that merging multiple assemblies and examining discordant regions using independent evidence is instrumental in assembling complex genomes.

Our biggest improvement to the assembly was on the Y chromosome, which has an unusual composition—its ~20 genes are interspersed among ~40 Mb of repetitive elements (Ritossa and Spiegelman 1965; Lohe *et al*. 1993; Carmena and Gonzalez 1995; Pimpinelli *et al*. 1995; Abad *et al*. 2004). Natural variation among *D. melanogaster* Y chromosomes can have broad effects on genome function and organismal fitness (*e.g*. Carvalho *et al*. 2000; Vibranovski *et al*. 2008; Paredes *et al*. 2011; Francisco and Lemos 2014; Kutch and Fedorka 2017; Wang *et al*. 2017). The extremely low nucleotide diversity of Y-linked genes (e.g. Zurovcova and Eanes 1999; Larracuente and Clark 2013; Morgan and Pardo-Manuel de Villena 2017) suggests that the Y-linked functional variation likely maps to the non-genic regions. The Y chromosome is a strong modifier of position effect variegation (PEV), a phenomenon that results in the stochastic silencing of euchromatic reporters caused by the spreading of heterochromatin (Karpen 1994; Elgin 1996; Wakimoto 1998). Y chromosomes may act as heterochromatin sinks, where extra Y-linked heterochromatin can titrate available heterochromatin-binding proteins away from other genomic locations. This may explain how genetic variation in Y-linked heterochromatin affects global gene expression (Henikoff 1996; Francisco and Lemos 2014; Brown and Bachtrog 2017). Alternatively, variation in Y-linked loci that generate small RNAs may have wide-scale impacts on chromatin organization (Zhou *et al*. 2012). These effects are difficult to tease apart without having a detailed view of Y chromosome sequence and organization. Our study discovered features of the Y chromosome that may relate to its interesting biology. Variation in Y-linked heterochromatin may affect the amount of silent chromatin marks in transposons (Brown and Bachtrog 2017), perhaps contributing to the higher rate of TE activity in males. We show that RNA transposons are generally overrepresented on the Y chromosome. It is possible that the overrepresentation of Y-linked retrotransposons is due to their increased activity in males: the Y chromosome heterochromatin sink effect may lead to reduced transcriptional silencing of TEs. In contrast to DNA transposons, the movement of retrotransposons is transcription dependent and therefore may result in differences in activity between the sexes. If the Y chromosome behaves as a sink for heterochromatin proteins, then we may expect the overrepresentation of RNA transposons to be a universal feature of Y chromosomes. Alternatively, differences in DNA repair or non-homologous recombination might lead to the differential accumulation of DNA and retrotransposons on the Y chromosome compared to the rest of the genome.

Y-linked structural variations can impact genome-wide gene regulatory variation in flies (*e.g. Su(Ste)* and *rDNA;* Lyckegaard and Clark 1989; Zhou *et al*. 2012) and male fertility in mammals (Reijo *et al*. 1995; Vogt *et al*. 1996; Sun *et al*. 2000; Repping *et al*. 2003; Morgan and Pardo-Manuel de Villena 2017). We find a large amount of gene traffic to the *D. melanogaster* Y chromosome from elsewhere in the genome. While estimates of interchromosomal duplications between the X and major autosomal arms range from ~3 (Bhutkar *et al*. 2007) to 7 (Han and Hahn 2012) on the *D. melanogaster* branch, we find at least 10 interchromosomal duplications to the Y chromosome. This observation is similar to other studies across taxa (Koerich *et al*. 2008; Hall *et al*. 2013; Hughes and Page 2015; Mahajan and Bachtrog 2017; Tobler *et al*. 2017). Our Y chromosome assembly provides new insights into the organization and mechanisms behind these duplications. For example, we found that most new translocations are DNA based and clustered in the Y pericentromic heterochromatin. The Y chromosome heterochromatin appears to be distinct from other heterochromatic regions of the genome, with properties that vary along the length of the chromosome (Wang *et al*. 2014). We hypothesize that the Y chromosome pericentromeric heterochromatin may be more accessible than other regions of the chromosome. If so, the increased accessibility may affect transcriptional activity and make these regions more prone to double stranded breaks (DSBs) that would facilitate structural rearrangements. Therefore, Y-linked pericentromeric chromatin may be more permissive to transcription compared to the rest of the chromosome allowing for natural selection to retain insertions that result in functional products. This may provide insights into how new Y-linked genes gain testis-specific functions. Notably, most Y-linked translocations are DNA-based and therefore involve DSB repair. Without a homolog to provide a template for DSB repair, microhomology-mediated end joining of non-homologous sequences may lead to insertions in the Y chromosome. DSB repair may also result in tandem duplications that contribute to the observed copy number variation in Y-linked genes. We discovered that most of the recent translocations to the Y chromosome exist in multiple copies (Table 2), suggesting that the tandem duplication rate may also be higher in the pericentric regions. However, most of these newly acquired genes are pseudogenized and are likely not constrained by natural selection. Many of the functional Y-linked genes are at least partially duplicated. Most essential Y-linked genes (*kl-2, kl-3, kl-5* and *ORY)* have larger introns (> 100Kb), with some introns reaching megabases in size (Kurek *et al*. 2000; Reugels *et al*. 2000). For genes with large overall sizes, complete gene duplications are less likely. In contrast, some functional genes, *e.g. rDNA, Mst77-Y* and Su(Ste), exist in multiple copies and are sensitive to gene dosage (Lyckegaard and Clark 1989; Zhou *et al*. 2012; Kost *et al*. 2015). A high duplication rate on the Y chromosome may therefore facilitate the evolution of Y-linked gene expression.

In mammals, some Y-linked genes have amplified into tandem arrays and exist in large palindromes (*e.g*. Rozen *et al*. 2003; Hughes *et al*. 2012; Soh *et al*. 2014). Gene conversion within these palindromes may be important for increasing the efficacy of selection on an otherwise non-recombining chromosome (Charlesworth 2003; Rozen *et al*. 2003; Connallon and Clark 2010). Interestingly, the largest gene families in the *D. melanogaster* genome, outside of the rDNA and histone clusters, are the Y-linked genes *Su(Ste)* and *PCKR*. We inferred a higher rate of gene conversion in both *PCKR* and *Su(Ste)* than the rest of the genome, and similar to the rate observed in mammalian Y chromosome (Rozen *et al*. 2003). However, our estimates do not consider recent selection or amplification of *PCKR* and Su(Ste). The elevated Y-linked gene conversion rates may be a consequence of having more highly amplified gene families than other genomic locations. Alternatively, the Y chromosome may have evolved distinct patterns of mutation because it lacks a homolog —low copy number Y-linked genes also have relatively high rates of gene conversion in Drosophila (Kopp *et al*. 2006) and humans (Rozen *et al*. 2003). Gene conversion between members of Y-linked multi-copy gene families may counteract the accumulation of deleterious mutations through evolutionary processes such as Muller’s ratchet (reviewed in Charlesworth and Charlesworth 2000; Charlesworth 2003; Rozen *et al*. 2003; Connallon and Clark 2010). If so, then we might expect high gene conversion rates to be a feature common among Y chromosomes.

## Acknowledgements

We thank Dr. Kevin Wei for feedback on the manuscript, and the University of Rochester Center for Integrated Research Computing for access to computing cluster resources. We also thank Drs. Casey Bergman, Matt Hahn, and Tom Eickbush for helpful discussion. This work was supported by National Institutes of Health National Institute of General Medical Sciences grant R35 GM119515-01 to A.M.L.

## SUPPLEMENTARY METHODS

### Genome assembly and reconciliation

For the reconciliation process, we generated two heterochromatin and two whole genome *de novo* assemblies with Falcon and Canu, independently. For the whole genome *de novo* assemblies, we used Canu v1.2 on all genomic reads with the parameters “genomeSize=160m useGrid=false errorRate=0.035” (Canu 1 assembly) and Falcon v0.3 (Falcon 1 assembly; configuration file is Supplementary text 1). We also generated *de novo* assemblies from the heterochromatin-enriched reads (see Methods) with Canu v1.3 (Canu 2 assembly) and Falcon v0.5 (Falcon 2 assembly; Supplementary text 2 for configuration file). To determine the best parameters for the heterochromatin-enriched Canu 2 assembly, we experimented with assembly conditions by creating *de novo* assemblies for all combinations of bogart em and ee between 0.025 and 0.06 (step size 0.005) for both the default Canu parameters and our repeat-sensitive parameters (“genomeSize=30m stopOnReadQuality=false corMinCoverage=0 corOutCoverage=100 ovlMerSize=31”). The assembly parameters that maximized N50, produced the longest total assembly size, and the longest contig length was bogart em and ee = 0.045. We therefore chose this assembly to represent Canu 2 for subsequent reconciliation steps. For the Falcon 2 assembly, we made assemblies by varying the minimal overlap length in the string graph (fc_ovlp_to_graph min_len 1000 and 6000) and chose min_len 1000 to represent the Falcon 2 assembly. In the next steps, we combined our *de novo* total and heterochromatin-enriched assemblies with reference assemblies from Chakraborty *et al*. 2016 (ISO_merged assembly) and release 6 (Hoskins *et al*. 2015).

We corrected any assembly errors manually. Our manual curation was primarily in detecting misassemblies in genic and intergenic regions according to the gene order in R6 using 154 heterochromatic and telomeric genes as our BLAST reference. After each reconciliation step, we split contigs with incorrect gene structures or genes from different chromosomal arms, as these likely were inappropriately merged by quickmerge or assemblers (Chakraborty *et al*. 2016). We first reconciled Falcon 1 and Canu 2 using Canu 2 as the reference (Merged 1). Merged 1 was reconciled with Falcon 2 using Merged 1 as the reference (Merged 2). We combined Merged 2 with the major chromosome arms in R6 (2L, 2R, 3L, 3R, 4, and X) using `cat` to create the Merged 3 assembly. To fill the gaps in Merged 3, we reconciled Merged 3 and ISO_merged (Chakraborty *et al*. 2016). using Merged 3 as the reference (Merged 4). Finally, the Merged 4 was reconciled with Canu 1 using Merged 4 as the reference (Final Merged). We corrected remaining assembly errors in the Final Merged assembly base on BLAST results and previous studies (see Methods). We polished the resulting final assembly with quiver and Pilon and used this version of the assembly for all subsequent analyses. We determined the order for the reconciliation process by: *1)* using combinations that improved the contiguity while retaining completeness; *2)* avoiding large-scale misassemblies due to the reconciliation process; and *3)* the ability to fill gaps (*e.g*. Canu 1 was useful for filling some gaps left in Merged 4).

### Estimating Y-linked gene conversion rates

Because Y-linked gene families do not undergo crossing over, we expect gene conversion to be the primary mechanism homogenizing different gene copies. We assume that there are a total of *n* copies of a gene, where *x* genes have the variant site that differentiates the copies, and for simplicity, any of the *n*-1 gene copies can convert a gene with equal probability. We also assume that there is no change in copy number. The fraction of differences between two gene copies at any generation *n* is given by *d_n_*.

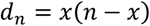

The effect of each gene conversion event will happen between copies with different SNPs or without SNPs. After the gene conversion, the divergence will be

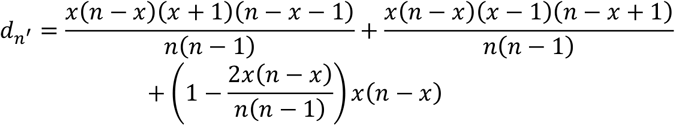

We can calculate the expected effect of each gene conversion on divergence.

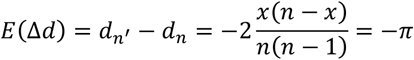

We assume parameter *c* is the rate at which a pair of gene copies homogenize each | other per generation, and corresponds to Ohta’s α (Ohta 1982). The divergence between copies is originated from point mutation with rate, u. If the divergence of gene family is only affected by gene conversion and mutations and the current divergence is under the gene conversion and mutation balance, we can derive,

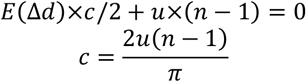

We can show that equation is equivalent to Rozen’s equation (Rozen *et al*. 2003) when *n* =2 and Ohta’s equation (Ohta 1982).

Here *c* is the rate of homogenized effect between 2 sequences by gene conversions. This rate is twice the rate that gene conversion happens. In addition, we need to consider the gene conversion tract length—we assumed that Y chromosome has the similar gene conversion tract length as other *D. melanogaster* chromosomes and normalize *c* based on 400 bp tract length of a single event (*c_g_*) (Miller *et al*. 2012; Miller *et al*. 2016).

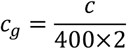

## Supplementary text 1. Falcon 1 configuration

[General]

input_fofn = input.fofn

input_type = raw

length_cutoff = 5000

length_cutoff_pr = 5000

jobqueue = production

job_type = local

sge_option_da = -pe smp 8 -q %(jobqueue)s

sge_option_la = -pe smp 2 -q %(jobqueue)s

sge_option_pda = -pe smp 8 -q %(jobqueue)s

sge_option_pla = -pe smp 2 -q %(jobqueue)s

sge_option_fc = -pe smp 24 -q %(jobqueue)s

sge_option_cns = -pe smp 8 -q %(jobqueue)s

pa_concurrent_jobs = 8

ovlp_concurrent_jobs = 8

pa_HPCdaligner_option = -v -dal128 -t8 -e.70 -l1000 -s1000 -M16

ovlp_HPCdaligner_option = -v -dal128 -t8 -h60 -e.96 -l500 -s1000 -M16

pa_DBsplit_option = -x500 -s400

ovlp_DBsplit_option = -x500 -s400

falcon_sense_option = --output_multi --min_idt 0.70 --min_cov 4 --

local_match_count_threshold 2 --max_n_read 200 --n_core 8 --output_dformatq overlap_filtering_setting = --max_diff 100 --max_cov 100 --min_cov 1 --bestn 10 -- n_core 8

## Supplementary text 2. Falcon 2 configuration

[General]

input_fofn = input_fal.fofn

input_type = raw

length_cutoff = -1

seed_coverage = 50

genome_size = 15000000

length_cutoff_pr = 1000

jobqueue = production

job_type = local

sge_option_da = -pe smp 8 -q %(jobqueue)s

sge_option_la = -pe smp 2 -q %(jobqueue)s

sge_option_pda = -pe smp 8 -q %(jobqueue)s

sge_option_pla = -pe smp 2 -q %(jobqueue)s

sge_option_fc = -pe smp 24 -q %(jobqueue)s

sge_option_cns = -pe smp 8 -q %(jobqueue)s

pa_concurrent_jobs = 9

ovlp_concurrent_jobs = 9

pa_HPCdaligner_option = -v -dal128 -t20 -H15000 -e.70 -k18 -w8 -l1000 -s100 - M24 -b

ovlp_HPCdaligner_option = -v -dal128 -t40 -M24 -k24 -h60 -e.95 -l500 -s100 - H15000 -b

pa_DBsplit_option = -x500 -s400

ovlp_DBsplit_option = -x500 -s400

falcon_sense_option = --output_multi --min_idt 0.70 --min_cov 1 --max_n_read 200 -- n_core 12

overlap_filtering_setting = --max_diff 100 --max_cov 100 --min_cov 1 --bestn 10 -- n_core 12

---

**Figure S1.**
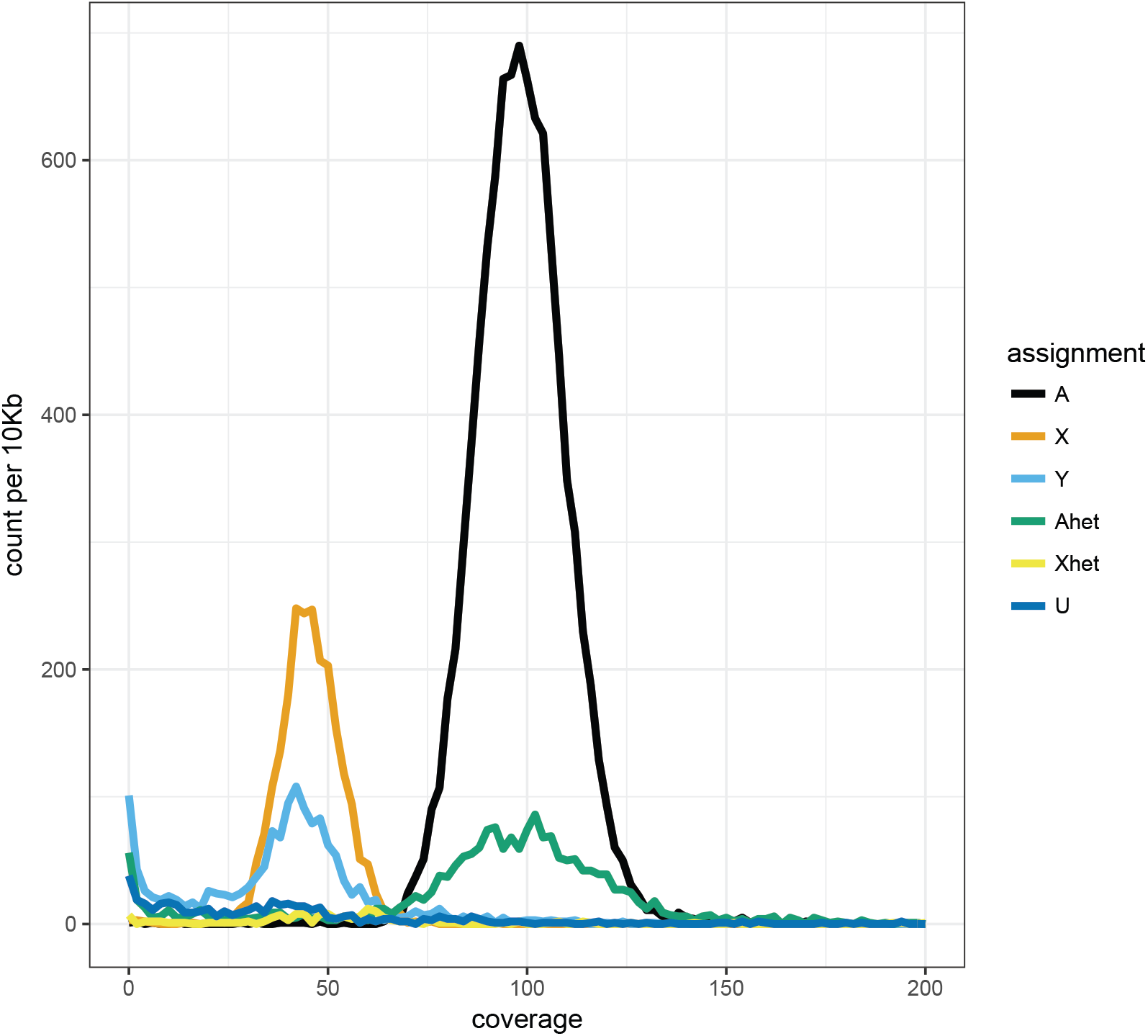
The median PacBio read coverage in different regions of genome. The raw Pacbio reads were mapped to our genome using minimap v2.5. We calculated median coverage of uniquely mapped reads using samtools and custom scripts. We assigned contig location by calculating the female-to-male coverage ratio of Illumina reads (see Methods). The heterochromatic autosomal and X chromosomal contigs were defined as contigs outside of the major contig for each chromosomal arm with the known chromosomal location. The raw data are available in Table S3.

**Figure S2.**
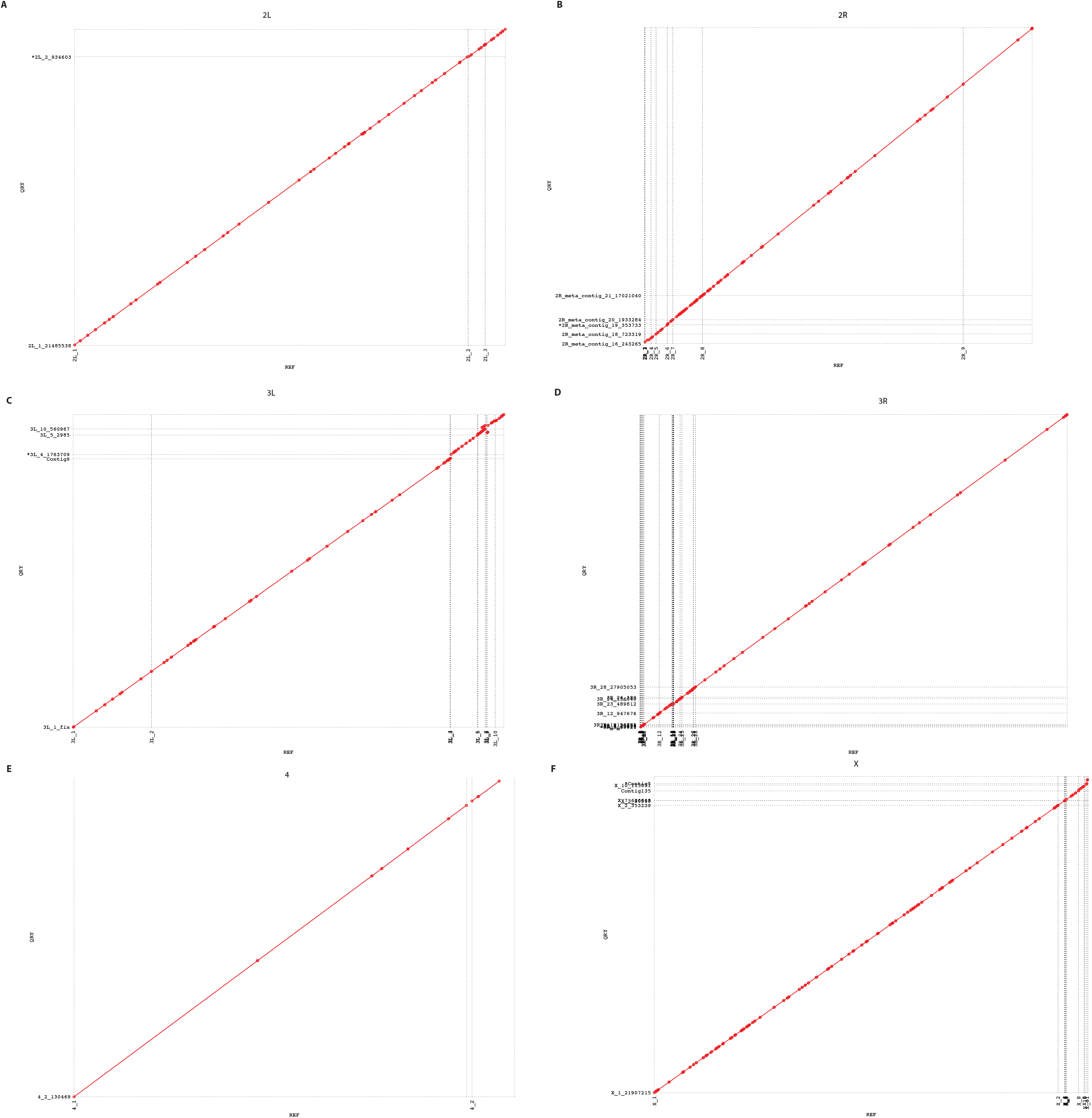
MUMMER plots showing whole genome alignments between the R6 assembly and our new assembly for autosomes and X chromosome. We mapped the contigs from our new assembly (y-axis) to the R6 cor (x-axis) using MUMMER, and only report one-to-one alignments.

**Figure S3.**
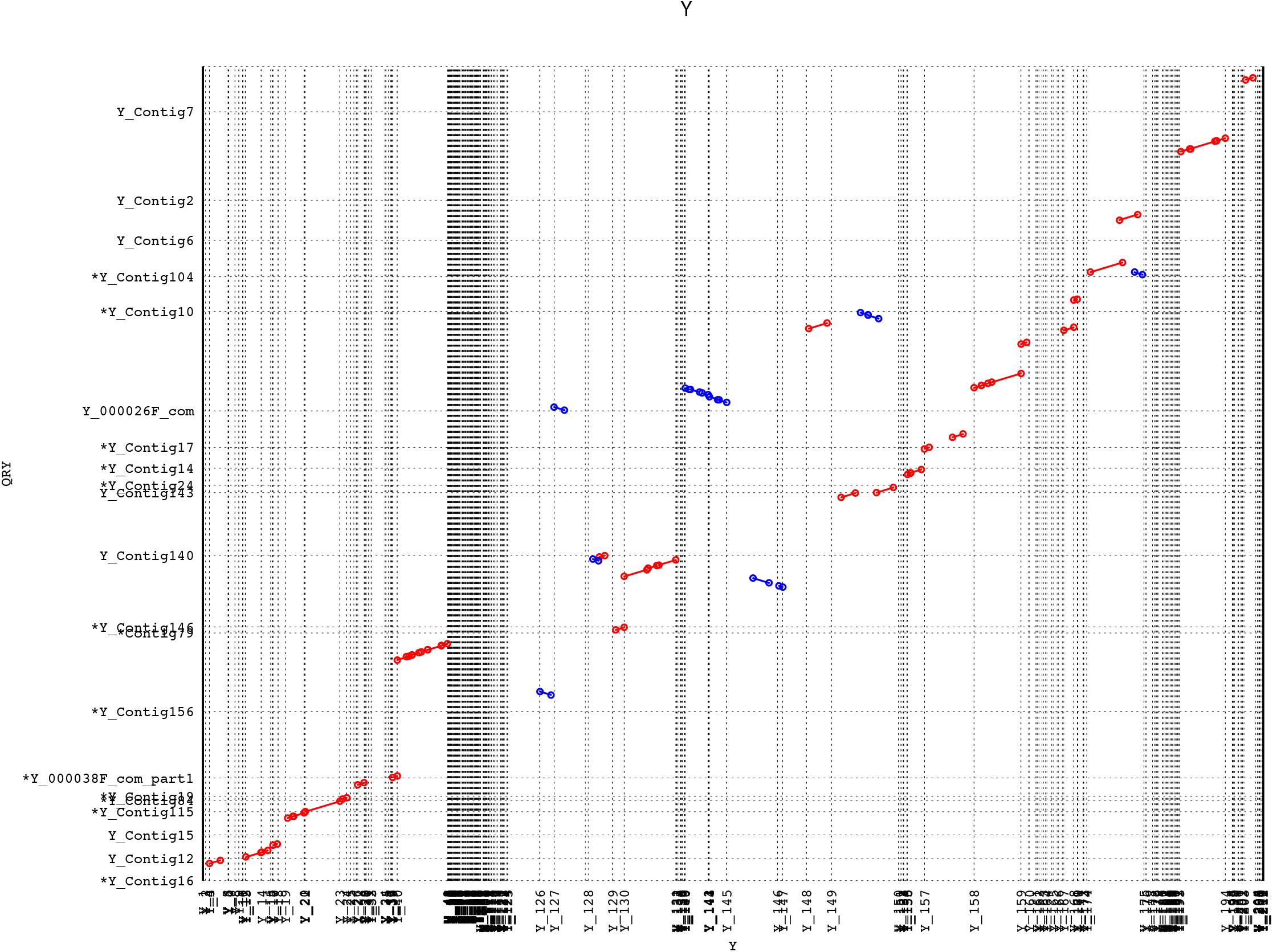
MUMMER plots showing the alignment between R6 Y chromosome assembly and our new Y chromosome assembly. We mapped the Y-linked contigs from our assembly (y-axis) to the R6 Y-linked contigs (x-axis) using MUMMER, and only report one-to-one alignments.

**Figure S4.**
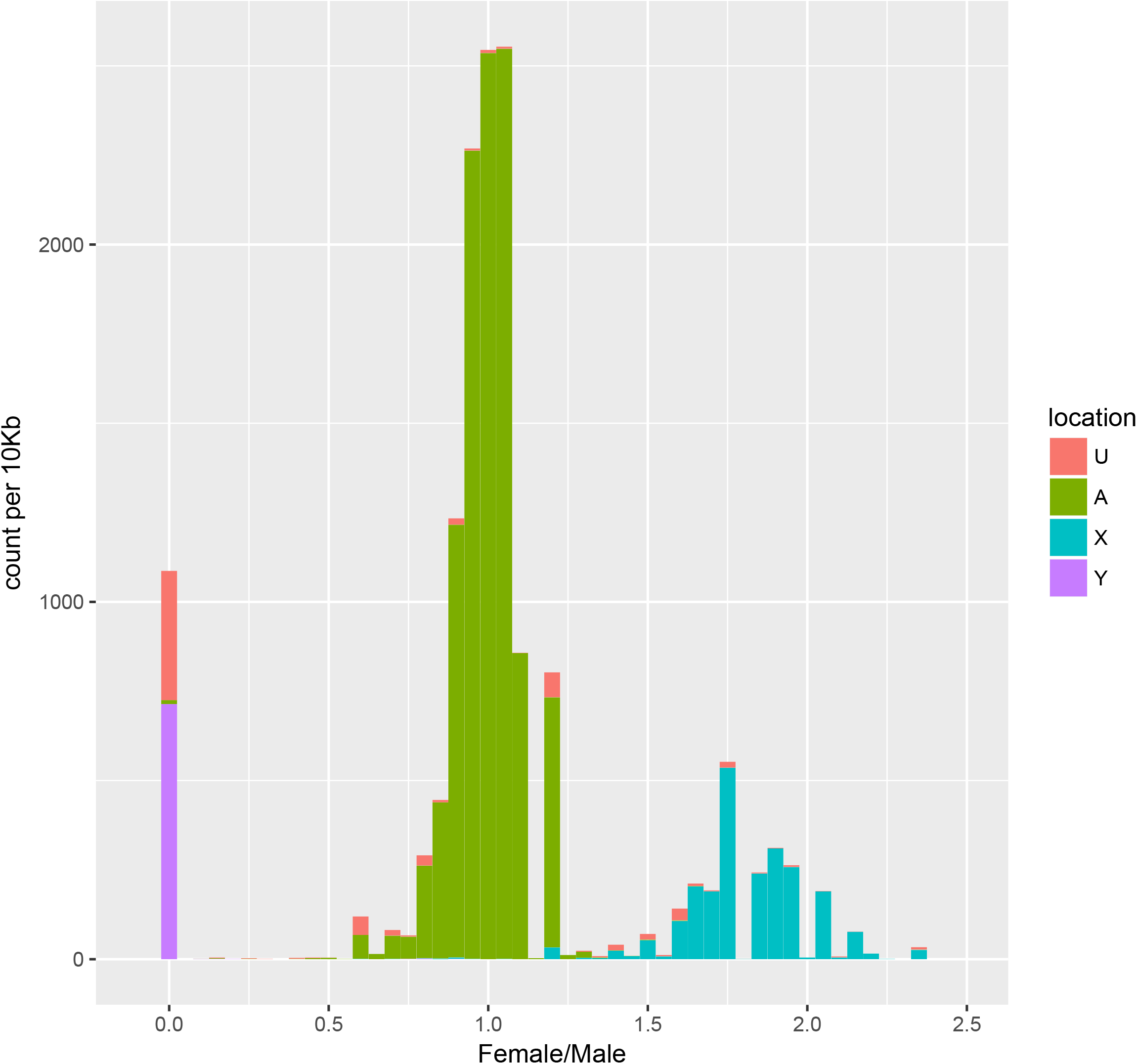
The median female-to-male coverage ratio of Illumina reads across different chromosomes based on the R6 annotation. We mapped the male and female Illumina reads to our new genome using bwa, and called median of female-to-male coverage ratio using samtools and custom scripts for each 10-kb region. The median female-to-male mapping ratio was normalized by total mapped reads. Contig location was determined by known gene content. Regions from contigs with Y-linked genes have a median female-to-male coverage ratio of 0.

**Figure S5.**
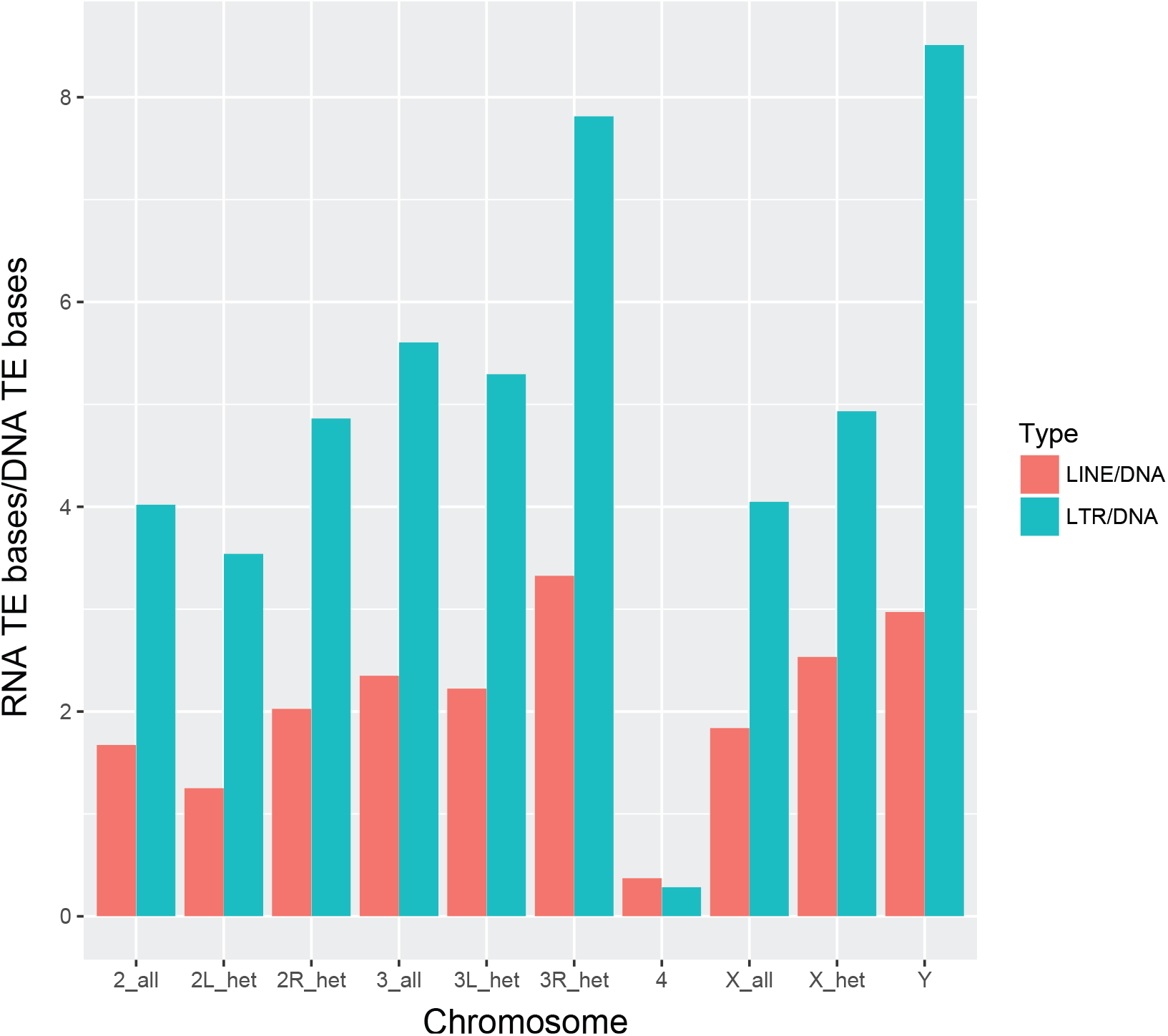
The repeat component in heterochromatic regions. We separated the repeat content by the chromatin states based on the coordinates in Hoskins et al. (2015). The total bases of LTR and LINE are normalized by the total bases of DNA transposons.

## REFERENCES

Abad, J. P., B. de Pablos, M. Agudo, I. Molina, G. Giovinazzo et al., 2004 Genomic and cytological analysis of the Y chromosome of *Drosophila melanogaster:* telomere-derived sequences at internal regions. Chromosoma 113: 295–304.

Agudo, M., A. Losada, J. P. Abad, S. Pimpinelli, P. Ripoll et al., 1999 Centromeres from telomeres? The centromeric region of the Y chromosome of *Drosophila melanogaster* contains a tandem array of telomeric *HeT-A-* and *TART*-related sequences. Nucleic Acids Res 27: 3318–3324.

Altschul, S. F., W. Gish, W. Miller, E. W. Myers and D. J. Lipman, 1990 Basic local alignment search tool. J Mol Biol 215: 403–410.

Backstrom, N., H. Ceplitis, S. Berlin and H. Ellegren, 2005 Gene conversion drives the evolution of HINTW, an ampliconic gene on the female-specific avian W chromosome. Mol Biol Evol 22: 1992–1999.

Bailly-Bechet, M., A. Haudry and E. Lerat, 2014 “One code to find them all”. a perl tool to conveniently parse RepeatMasker output files. Mobile DNA 5: 13.

Balakireva, M. D., Y. Shevelyov, D. I. Nurminsky, K. J. Livak and V. A. Gvozdev, 1992 Structural organization and diversification of Y-linked sequences comprising *Su(Ste)* genes in *Drosophila melanogaster.* Nucleic Acids Res 20: 3731–3736.

Bayes, J. J., and H. S. Malik, 2009 Altered heterochromatin binding by a hybrid sterility protein in Drosophila sibling species. Science 326: 1538–1541.

Benson, G., 1999 Tandem repeats finder. a program to analyze DNA sequences. Nucleic Acids Res 27: 573–580.

Bhutkar, A., W. M. Gelbart and T. F. Smith, 2007 Inferring genome-scale rearrangement phylogeny and ancestral gene order. a Drosophila case study. Genome Biol 8: R236.

Bonaccorsi, S., and A. Lohe, 1991 Fine mapping of satellite DNA sequences along the Y chromosome of *Drosophila melanogaster:* relationships between satellite sequences and fertility factors. Genetics 129: 177–189.

Bozzetti, M. P., S. Massari, P. Finelli, F. Meggio, L. A. Pinna et al., 1995 The *Ste* locus, a component of the parasitic *cry-Ste* system of *Drosophila melanogaster*, encodes a protein that forms crystals in primary spermatocytes and mimics properties of the beta subunit of casein kinase 2. Proc Natl Acad Sci U S A 92: 6067–6071.

Branton, D., D. W. Deamer, A. Marziali, H. Bayley, S. A. Benner et al., 2008 The potential and challenges of nanopore sequencing. Nat Biotechnol 26: 11461153.

Brown, E., and D. Bachtrog, 2017 The Drosophila Y chromosome affects heterochromatin integrity genome-wide. bioRxiv.

Brown, E. J., and D. Bachtrog, 2014 The chromatin landscape of Drosophila. comparisons between species, sexes, and chromosomes. Genome Res 24: 1125–1137.

Bucheton, A., J. M. Lavige, G. Picard and P. L’Heritier, 1976 Non-mendelian female sterility in *Drosophila melanogaster.* quantitative variations in the efficiency of inducer and reactive strains. Heredity (Edinb) 36: 305–314.

Carlson, M., and D. Brutlag, 1977 Cloning and characterization of a complex satellite DNA from *Drosophila melanogaster.* Cell 11: 371–381.

Carmena, M., and C. Gonzalez, 1995 Transposable elements map in a conserved pattern of distribution extending from beta-heterochromatin to centromeres in *Drosophila melanogaster.* Chromosoma 103: 676–684.

Carvalho, A. B., B. A. Dobo, M. D. Vibranovski and A. G. Clark, 2001 Identification of five new genes on the Y chromosome of *Drosophila melanogaster.* Proc Natl Acad Sci U S A 98: 13225–13230.

Carvalho, A. B., E. G. Dupim and G. Goldstein, 2016 Improved assembly of noisy long reads by k-mer validation. Genome Res 26: 1710–1720.

Carvalho, A. B., B. P. Lazzaro and A. G. Clark, 2000 Y chromosomal fertility factors *kl-2* and *kl-3* of *Drosophila melanogaster* encode dynein heavy chain polypeptides. Proc Natl Acad Sci U S A 97: 13239–13244.

Carvalho, A. B., M. D. Vibranovski, J. W. Carlson, S. E. Celniker, R. A. Hoskins et al., 2003 Y chromosome and other heterochromatic sequences of the *Drosophila melanogaster* genome: how far can we go? Genetica 117: 227–237.

Carvalho, A. B., B. Vicoso, C. A. Russo, B. Swenor and A. G. Clark, 2015 Birth of a new gene on the Y chromosome of *Drosophila melanogaster.* Proc Natl Acad Sci U S A 112: 12450–12455.

Cattani, M. V., and D. C. Presgraves, 2012 Incompatibility between X chromosome factor and pericentric heterochromatic region causes lethality in hybrids between *Drosophila melanogaster* and its sibling species. Genetics 191: 549–559.

Chaisson, M. J., J. Huddleston, M. Y. Dennis, P. H. Sudmant, M. Malig et al., 2015 Resolving the complexity of the human genome using single-molecule sequencing. Nature 517: 608–611.

Chaisson, M. J., and G. Tesler, 2012 Mapping single molecule sequencing reads using basic local alignment with successive refinement (BLASR): application and theory. BMC Bioinformatics 13: 238.

Chakraborty, M., J. G. Baldwin-Brown, A. D. Long and J. J. Emerson, 2016 Contiguous and accurate de novo assembly of metazoan genomes with modest long read coverage. Nucleic Acids Res 44: e147.

Chang, C. H., and A. M. Larracuente, 2017 Genomic changes following the reversal of a Y chromosome to an autosome in *Drosophila pseudoobscura.* Evolution 71: 1285–1296.

Charlesworth, B., 2003 The organization and evolution of the human Y chromosome. Genome Biol 4: 226.

Charlesworth, B., and D. Charlesworth, 2000 The degeneration of Y chromosomes. Philos Trans R Soc Lond B Biol Sci 355: 1563–1572.

Charlesworth, B., C. H. Langley and W. Stephan, 1986 The evolution of restricted recombination and the accumulation of repeated DNA sequences. Genetics 112: 947–962.

Chin, C. S., D. H. Alexander, P. Marks, A. A. Klammer, J. Drake et al., 2013 Nonhybrid, finished microbial genome assemblies from long-read SMRT sequencing data. Nat Methods 10: 563–569.

Chin, C. S., P. Peluso, F. J. Sedlazeck, M. Nattestad, G. T. Concepcion et al., 2016 Phased diploid genome assembly with single-molecule real-time sequencing. Nat Methods 13: 1050–1054.

Chippindale, A. K., and W. R. Rice, 2001 Y chromosome polymorphism is a strong determinant of male fitness in *Drosophila melanogaster.* Proc Natl Acad Sci U S A 98: 5677–5682.

Clifton, B. D., P. Librado, S. D. Yeh, E. S. Solares, D. A. Real et al., 2017 Rapid Functional and Sequence Differentiation of a Tandemly Repeated Species-Specific Multigene Family in Drosophila. Mol Biol Evol 34: 51–65.

Comeron, J. M., R. Ratnappan and S. Bailin, 2012 The many landscapes of recombination in *Drosophila melanogaster.* PLoS Genet 8: e1002905.

Connallon, T., and A. G. Clark, 2010 Gene duplication, gene conversion and the evolution of the Y chromosome. Genetics 186: 277–286.

Csink, A. K., and S. Henikoff, 1996 Genetic modification of heterochromatic association and nuclear organization in Drosophila. Nature 381: 529–531.

Danilevskaya, O. N., E. V. Kurenova, M. N. Pavlova, D. V. Bebehov, A. J. Link et al., 1991 He-T family DNA sequences in the Y chromosome of *Drosophila melanogaster* share homology with the X-linked stellate genes. Chromosoma 100: 118–124.

Dernburg, A. F., J. W. Sedat and R. S. Hawley, 1996 Direct evidence of a role for heterochromatin in meiotic chromosome segregation. Cell 86: 135–146.

Dimitri, P., and C. Pisano, 1989 Position effect variegation in *Drosophila melanogaster:* relationship between suppression effect and the amount of Y chromosome. Genetics 122: 793–800.

Eid, J., A. Fehr, J. Gray, K. Luong, J. Lyle et al., 2009 Real-time DNA sequencing from single polymerase molecules. Science 323: 133–138.

Elgin, S. C., 1996 Heterochromatin and gene regulation in Drosophila. Curr Opin Genet Dev 6: 193–202.

Ferree, P. M., and D. A. Barbash, 2009 Species-specific heterochromatin prevents mitotic chromosome segregation to cause hybrid lethality in Drosophila. PLoS Biol 7: e1000234.

Francisco, F. O., and B. Lemos, 2014 How do y-chromosomes modulate genome-wide epigenetic States. genome folding, chromatin sinks, and gene expression. J Genomics 2: 94–103.

Garavis, M., M. Mendez-Lago, V. Gabelica, S. L. Whitehead, C. Gonzalez et al., 2015 The structure of an endogenous Drosophila centromere reveals the prevalence of tandemly repeated sequences able to form i-motifs. Sci Rep 5: 13307.

Gatti, M., and S. Pimpinelli, 1992 Functional elements in *Drosophila melanogaster* heterochromatin. Annu Rev Genet 26: 239–275.

Hall, A. B., Y. Qi, V. Timoshevskiy, M. V. Sharakhova, I. V. Sharakhov et al., 2013 Six novel Y chromosome genes in Anopheles mosquitoes discovered by independently sequencing males and females. BMC Genomics 14: 273.

Han, M. V., and M. W. Hahn, 2012 Inferring the history of interchromosomal gene transposition in Drosophila using n-dimensional parsimony. Genetics 190: 813–825.

Henikoff, S., 1996 Dosage-dependent modification of position-effect variegation in Drosophila. Bioessays 18: 401–409.

Hoskins, R. A., J. W. Carlson, K. H. Wan, S. Park, I. Mendez et al., 2015 The Release 6 reference sequence of the *Drosophila melanogaster* genome. Genome Res 25: 445–458.

Hoskins, R. A., C. D. Smith, J. W. Carlson, A. B. Carvalho, A. Halpern et al., 2002 Heterochromatic sequences in a Drosophila whole-genome shotgun assembly. Genome Biol 3: RESEARCH0085.

Huddleston, J., S. Ranade, M. Malig, F. Antonacci, M. Chaisson et al., 2014 Reconstructing complex regions of genomes using long-read sequencing technology. Genome Res 24: 688–696.

Hudson, R. R., 1987 Estimating the recombination parameter of a finite population model without selection. Genet Res 50: 245–250.

Hughes, J. F., and D. C. Page, 2015 The Biology and Evolution of Mammalian Y Chromosomes. Annu Rev Genet 49: 507–527.

Hughes, J. F., H. Skaletsky, L. G. Brown, T. Pyntikova, T. Graves et al., 2012 Strict evolutionary conservation followed rapid gene loss on human and rhesus Y chromosomes. Nature 483: 82–86.

Hurst, L. D., 1996 Further evidence consistent with Stellate’s involvement in meiotic drive. Genetics 142: 641–643.

Jagannathan, M., N. Warsinger-Pepe, G. J. Watase and Y. M. Yamashita, 2017 Comparative Analysis of Satellite DNA in the *Drosophila melanogaster* Species Complex. G3 (Bethesda) 7: 693–704.

Jain, M., H. E. Olsen, D. J. Turner, D. Stoddart, K. V. Bulazel et al., 2017 Linear Assembly of a Human Y Centromere using Nanopore Long Reads. bioRxiv.

Junakovic, N., A. Terrinoni, C. Di Franco, C. Vieira and C. Loevenbruck, 1998 Accumulation of transposable elements in the heterochromatin and on the Y chromosome of Drosophila simulans and *Drosophila melanogaster.* J Mol Evol 46: 661–668.

Karpen, G. H., 1994 Position-effect variegation and the new biology of heterochromatin. Curr Opin Genet Dev 4: 281–291.

Kearse, M., R. Moir, A. Wilson, S. Stones-Havas, M. Cheung et al., 2012 Geneious Basic: an integrated and extendable desktop software platform for the organization and analysis of sequence data. Bioinformatics 28: 1647–1649.

Keightley, P. D., R. W. Ness, D. L. Halligan and P. R. Haddrill, 2014 Estimation of the spontaneous mutation rate per nucleotide site in a Drosophila melanogaster full-sib family. Genetics 196: 313–320.

Kennison, J. A., 1981 The Genetic and Cytological Organization of the Y Chromosome of DROSOPHILA MELANOGASTER. Genetics 98: 529–548.

Kent, W. J., 2002 BLAT--the BLAST-like alignment tool. Genome Res 12: 656–664.

Khost, D. E., D. G. Eickbush and A. M. Larracuente, 2017 Single-molecule sequencing resolves the detailed structure of complex satellite DNA loci in *Drosophila melanogaster.* Genome Res 27: 709–721.

Kim, D., B. Langmead and S. L. Salzberg, 2015 HISAT: a fast spliced aligner with low memory requirements. Nat Methods 12: 357–360.

Kim, K. E., P. Peluso, P. Babayan, P. J. Yeadon, C. Yu et al., 2014 Long-read, whole-genome shotgun sequence data for five model organisms. Sci Data 1: 140045.

Koerich, L. B., X. Wang, A. G. Clark and A. B. Carvalho, 2008 Low conservation of gene content in the Drosophila Y chromosome. Nature 456: 949–951.

Kogan, G. L., V. N. Epstein, A. A. Aravin and V. A. Gvozdev, 2000 Molecular evolution of two paralogous tandemly repeated heterochromatic gene clusters linked to the X and Y chromosomes of *Drosophila melanogaster.* Mol Biol Evol 17: 697–702.

Kogan, G. L., L. A. Usakin, S. S. Ryazansky and V. A. Gvozdev, 2012 Expansion and evolution of the X-linked testis specific multigene families in the melanogaster species subgroup. PLoS One 7: e37738.

Kopp, A., A. K. Frank and O. Barmina, 2006 Interspecific divergence, intrachromosomal recombination, and phylogenetic utility of Y-chromosomal genes in Drosophila. Mol Phylogenet Evol 38: 731–741.

Koren, S., B. P. Walenz, K. Berlin, J. R. Miller, N. H. Bergman et al., 2017 Canu. scalable and accurate long-read assembly via adaptive k-mer weighting and repeat separation. Genome Res 27: 722–736.

Kost, N., S. Kaiser, Y. Ostwal, D. Riedel, A. Stutzer et al., 2015 Multimerization of Drosophila sperm protein Mst77F causes a unique condensed chromatin structure. Nucleic Acids Res 43: 3033–3045.

Krsticevic, F. J., H. L. Santos, S. Januario, C. G. Schrago and A. B. Carvalho, 2010 Functional copies of the *Mst77F* gene on the Y chromosome of *Drosophila melanogaster.* Genetics 184: 295–307.

Krsticevic, F. J., C. G. Schrago and A. B. Carvalho, 2015 Long-Read Single Molecule Sequencing to Resolve Tandem Gene Copies. The Mst77Y Region on the *Drosophila melanogaster* Y Chromosome. G3 (Bethesda) 5: 1145–1150.

Kurek, R., A. M. Reugels, U. Lammermann and H. Bunemann, 2000 Molecular aspects of intron evolution in dynein encoding mega-genes on the heterochromatic Y chromosome of Drosophila sp. Genetica 109: 113–123.

Kurtz, S., A. Phillippy, A. L. Delcher, M. Smoot, M. Shumway et al., 2004 Versatile and open software for comparing large genomes. Genome Biol 5: R12.

Kutch, I. C., and K. M. Fedorka, 2017 A test for Y-linked additive and epistatic effects on surviving bacterial infections in *Drosophila melanogaster.* J Evol Biol 30: 1400–1408.

Larracuente, A. M., and A. G. Clark, 2013 Surprising differences in the variability of Y chromosomes in African and cosmopolitan populations of *Drosophila melanogaster.* Genetics 193: 201–214.

Lemos, B., L. O. Araripe and D. L. Hartl, 2008 Polymorphic Y chromosomes harbor cryptic variation with manifold functional consequences. Science 319: 91–93.

Lemos, B., A. T. Branco and D. L. Hartl, 2010 Epigenetic effects of polymorphic Y chromosomes modulate chromatin components, immune response, and sexual conflict. Proc Natl Acad Sci U S A 107: 15826–15831.

Li, H., 2016 Minimap and miniasm. fast mapping and de novo assembly for noisy long sequences. Bioinformatics 32: 2103–2110.

Li, H., and R. Durbin, 2010 Fast and accurate long-read alignment with BurrowsWheeler transform. Bioinformatics 26: 589–595.

Li, H., B. Handsaker, A. Wysoker, T. Fennell, J. Ruan et al., 2009 The Sequence Alignment/Map format and SAMtools. Bioinformatics 25: 2078–2079.

Livak, K. J., 1984 Organization and mapping of a sequence on the *Drosophila melanogaster* X and Y chromosomes that is transcribed during spermatogenesis. Genetics 107: 611–634.

Lohe, A. R., and D. L. Brutlag, 1987a Adjacent satellite DNA segments in Drosophila structure of junctions. J Mol Biol 194: 171–179.

Lohe, A. R., and D. L. Brutlag, 1987b Identical satellite DNA sequences in sibling species of Drosophila. J Mol Biol 194: 161–170.

Lohe, A. R., A. J. Hilliker and P. A. Roberts, 1993 Mapping simple repeated DNA sequences in heterochromatin of *Drosophila melanogaster.* Genetics 134: 1149–1174.

Lyckegaard, E. M., and A. G. Clark, 1989 Ribosomal DNA and Stellate gene copy number variation on the Y chromosome of *Drosophila melanogaster.* Proc Natl Acad Sci U S A 86: 1944–1948.

Mahajan, S., and D. Bachtrog, 2017 Convergent evolution of Y chromosome gene content in flies. Nat Commun 8: 785.

Mahajan, S., K. Wei, M. Nalley, L. Gibilisco and D. Bachtrog, 2018.

Malone, C. D., R. Lehmann and F. K. Teixeira, 2015 The cellular basis of hybrid dysgenesis and Stellate regulation in Drosophila. Curr Opin Genet Dev 34: 88–94.

McKee, B. D., C. S. Hong and S. Das, 2000 On the roles of heterochromatin and euchromatin in meiosis in drosophila: mapping chromosomal pairing sites and testing candidate mutations for effects on X-Y nondisjunction and meiotic drive in male meiosis. Genetica 109: 77–93.

McKee, B. D., and M. T. Satter, 1996 Structure of the Y chromosomal *Su(Ste)* locus in *Drosophila melanogaster* and evidence for localized recombination among repeats. Genetics 142: 149–161.

Mendez-Lago, M., C. M. Bergman, B. de Pablos, A. Tracey, S. L. Whitehead et al., 2011 A large palindrome with interchromosomal gene duplications in the pericentromeric region of the *D. melanogaster* Y chromosome. Mol Biol Evol 28: 1967–1971.

Mendez-Lago, M., J. Wild, S. L. Whitehead, A. Tracey, B. de Pablos et al., 2009 Novel sequencing strategy for repetitive DNA in a Drosophila BAC clone reveals that the centromeric region of the Y chromosome evolved from a telomere. Nucleic Acids Res 37: 2264–2273.

Miller, D. E., C. B. Smith, N. Y. Kazemi, A. J. Cockrell, A. V. Arvanitakas et al., 2016 Whole-Genome Analysis of Individual Meiotic Events in Drosophila melanogaster Reveals That Noncrossover Gene Conversions Are Insensitive to Interference and the Centromere Effect. Genetics 203: 159–171.

Miller, D. E., S. Takeo, K. Nandanan, A. Paulson, M. M. Gogol et al., 2012 A Whole-Chromosome Analysis of Meiotic Recombination in *Drosophila melanogaster.* G3 (Bethesda) 2: 249–260.

Morgan, A. P., and F. Pardo-Manuel de Villena, 2017 Sequence and Structural Diversity of Mouse Y Chromosomes. Mol Biol Evol 34: 3186–3204.

Nishida, K. M., K. Saito, T. Mori, Y. Kawamura, T. Nagami-Okada et al., 2007 Gene silencing mechanisms mediated by Aubergine piRNA complexes in Drosophila male gonad. RNA 13: 1911–1922.

Nurminsky, D. I., Y. Shevelyov, S. V. Nuzhdin and V. A. Gvozdev, 1994 Structure, molecular evolution and maintenance of copy number of extended repeated structures in the X-heterochromatin of *Drosophila melanogaster.* Chromosoma 103: 277–285.

Ohta, T., 1984 Some models of gene conversion for treating the evolution of multigene families. Genetics 106: 517–528.

Paradis, E., J. Claude and K. Strimmer, 2004 APE: Analyses of Phylogenetics and Evolution in R language. Bioinformatics 20: 289–290.

Paredes, S., A. T. Branco, D. L. Hartl, K. A. Maggert and B. Lemos, 2011 Ribosomal DNA deletions modulate genome-wide gene expression: “rDNA-sensitive” genes and natural variation. PLoS Genet 7: e1001376.

Pertea, M., G. M. Pertea, C. M. Antonescu, T. C. Chang, J. T. Mendell et al., 2015 StringTie enables improved reconstruction of a transcriptome from RNA-seq reads. Nat Biotechnol 33: 290–295.

Pimpinelli, S., M. Berloco, L. Fanti, P. Dimitri, S. Bonaccorsi et al., 1995 Transposable elements are stable structural components of *Drosophila melanogaster* heterochromatin. Proc Natl Acad Sci U S A 92: 3804–3808.

Reijo, R., T. Y. Lee, P. Salo, R. Alagappan, L. G. Brown et al., 1995 Diverse spermatogenic defects in humans caused by Y chromosome deletions encompassing a novel RNA-binding protein gene. Nat Genet 10: 383–393.

Repping, S., H. Skaletsky, L. Brown, S. K. van Daalen, C. M. Korver et al., 2003 Polymorphism for a 1.6-Mb deletion of the human Y chromosome persists through balance between recurrent mutation and haploid selection. Nat Genet 35: 247–251.

Reugels, A. M., R. Kurek, U. Lammermann and H. Bunemann, 2000 Mega-introns in the dynein gene DhDhc7(Y) on the heterochromatic Y chromosome give rise to the giant threads loops in primary spermatocytes of Drosophila hydei. Genetics 154: 759–769.

Ritossa, F. M., and S. Spiegelman, 1965 Localization of DNA Complementary to Ribosomal Rna in the Nucleolus Organizer Region of *Drosophila melanogaster.* Proc Natl Acad Sci U S A 53: 737–745.

Rohmer, C., J. R. David, B. Moreteau and D. Joly, 2004 Heat induced male sterility in *Drosophila melanogaster*: adaptive genetic variations among geographic populations and role of the Y chromosome. J Exp Biol 207: 2735–2743.

Rosic, S., F. Kohler and S. Erhardt, 2014 Repetitive centromeric satellite RNA is essential for kinetochore formation and cell division. J Cell Biol 207: 335–349.

Rozen, S., H. Skaletsky, J. D. Marszalek, P. J. Minx, H. S. Cordum et al., 2003 Abundant gene conversion between arms of palindromes in human and ape Y chromosomes. Nature 423: 873–876.

Shevelyov, Y. Y., 1992 Copies of a Stellate gene variant are located in the X heterochromatin of *Drosophila melanogaster* and are probably expressed. Genetics 132: 1033–1037.

Smit, A., R. Hubley and P. Green, 2013 RepeatMasker, pp. in Open-4.0.

Soh, Y. Q., J. Alfoldi, T. Pyntikova, L. G. Brown, T. Graves et al., 2014 Sequencing the mouse Y chromosome reveals convergent gene acquisition and amplification on both sex chromosomes. Cell 159: 800–813.

Sun, C., H. Skaletsky, S. Rozen, J. Gromoll, E. Nieschlag et al., 2000 Deletion of azoospermia factor a (AZFa) region of human Y chromosome caused by recombination between HERV15 proviruses. Hum Mol Genet 9: 2291–2296.

Thornton, K., 2003 Libsequence. a C++ class library for evolutionary genetic analysis. Bioinformatics 19: 2325–2327.

Tobler, R., V. Nolte and C. Schlotterer, 2017 High rate of translocation-based gene birth on the Drosophila Y chromosome. Proc Natl Acad Sci U S A 114: 11721–11726.

Traverse, K. L., and M. L. Pardue, 1989 Studies of He-T DNA sequences in the pericentric regions of Drosophila chromosomes. Chromosoma 97: 261–271.

Treangen, T. J., and S. L. Salzberg, 2011 Repetitive DNA and next-generation sequencing. computational challenges and solutions. Nat Rev Genet 13: 36–46.

Tulin, A. V., G. L. Kogan, D. Filipp, M. D. Balakireva and V. A. Gvozdev, 1997 Heterochromatic Stellate gene cluster in *Drosophila melanogaster*. structure and molecular evolution. Genetics 146: 253–262.

Usakin, L. A., G. L. Kogan, A. I. Kalmykova and V. A. Gvozdev, 2005 An alien promoter capture as a primary step of the evolution of testes-expressed repeats in the *Drosophila melanogaster* genome. Mol Biol Evol 22: 1555–1560.

Vibranovski, M. D., L. B. Koerich and A. B. Carvalho, 2008 Two new Y-linked genes in *Drosophila melanogaster.* Genetics 179: 2325–2327.

Vogt, P. H., A. Edelmann, S. Kirsch, O. Henegariu, P. Hirschmann et al., 1996 Human Y chromosome azoospermia factors (AZF) mapped to different subregions in Yq11. Hum Mol Genet 5: 933–943.

Wakimoto, B. T., 1998 Beyond the nucleosome. epigenetic aspects of position-effect variegation in Drosophila. Cell 93: 321–324.

Walker, B. J., T. Abeel, T. Shea, M. Priest, A. Abouelliel et al., 2014 Pilon. an integrated tool for comprehensive microbial variant detection and genome assembly improvement. PLoS One 9: e112963.

Wang, M., A. T. Branco and B. Lemos, 2017 The Y Chromosome Modulates Splicing and Sex-Biased Intron Retention Rates in Drosophila. Genetics.

Wang, S. H., R. Nan, M. C. Accardo, M. Sentmanat, P. Dimitri et al., 2014 A distinct type of heterochromatin at the telomeric region of the Drosophila melanogaster Y chromosome. PLoS One 9: e86451.

Wei, K. H. C., S. E. Lower, I. V. Caldas, T. J. Sless, D. A. Barbash et al., 2018 Variable rates of simple satellite gains across the Drosophila phylogeny. Molecular Biology and Evolution: msy005–msy005.

Zhao, H., Z. Sun, J. Wang, H. Huang, J. P. Kocher et al., 2014 CrossMap. a versatile tool for coordinate conversion between genome assemblies. Bioinformatics 30: 1006–1007.

Zhou, J., T. B. Sackton, L. Martinsen, B. Lemos, T. H. Eickbush et al., 2012 Y chromosome mediates ribosomal DNA silencing and modulates the chromatin state in Drosophila. Proc Natl Acad Sci U S A 109: 9941–9946.

Zimin, A. V., D. Puiu, M. C. Luo, T. Zhu, S. Koren et al., 2017 Hybrid assembly of the large and highly repetitive genome of *Aegilops tauschii*, a progenitor of bread wheat, with the MaSuRCA mega-reads algorithm. Genome Res 27: 787–792.

Zurovcova, M., and W. F. Eanes, 1999 Lack of nucleotide polymorphism in the Y-linked sperm flagellar dynein gene Dhc-Yh3 of *Drosophila melanogaster* and D. simulans. Genetics 153: 1709–1715.

## REFERENCES

Ohta, T., 1982 Allelic and nonallelic homology of a supergene family. Proc Natl Acad Sci U S A 79: 3251–3254.

